# How does housing quality affect maternal investment and reproduction?

**DOI:** 10.64898/2026.10.17.759638

**Authors:** Prathipa Anandarajan, Georgia Mason

## Abstract

Poor housing is linked to reproductive failure in captive mammals, but the mechanisms are rarely tested. To understand the mechanism behind it, we used DBA/2 mice as models and in conventional housing (CH) and well-resourced housing (WR) across three cohorts (N = 36 sister-pair cages; 6 CH and 6 WR per Cohort). We tested two mutually non-exclusive mechanisms: the energy hypothesis (low maternal body condition, measured as BMI, impairs reproduction) and the stereotypic behaviour (SB) hypothesis (SB competes with reproduction and maternal care regardless of body condition). Housing affected maternal care: WR dams built better nests (p = 0.002), nursed more (p = 0.018) and nursed for more days before weaning (p = 0.023), and their pups tended to be heavier at PND 28 (14.2 vs 12.5 g, p = 0.06). Unexpectedly, all CH dams (7/7) retrieved displaced pups, but no WR dams did (0/8), likely reflecting how the test cage was perceived. But housing did not affect mating, pregnancy, litter size, pup survival or pup weight up to postnatal day (PND) 21. CH females performed about 9 times more SB than WR females before mating, and the difference persisted through lactation. BMI did not differ by housing, so the energy hypothesis was not supported. Adding SB to models weakened housing effects on nest quality and nursing, but SB itself did not significantly predict any outcome. Here, we found that housing quality mainly shapes how dams care for pups during lactation but not their capacity to breed. SB may play a partial role, but that needs better-powered testing. This study supports improving housing for captive breeding programmes now, even though the mechanism remains unknown.

## 1. Introduction

Housing quality can affect nearly every stage of mammalian reproduction, from mate choice to weaning, and sub-optimal housing tends to make outcomes worse. At least six mechanisms could explain this: neuroendocrine dysregulation, depression-like anhedonia, excessive maternal weight gain, excessive maternal weight loss, abnormal repetitive behaviour (ARB), and reduced mate quality altering female investment. Here we test two of them: maternal weight loss (the energy hypothesis) and ARB (Table 1).

**Table 1.**
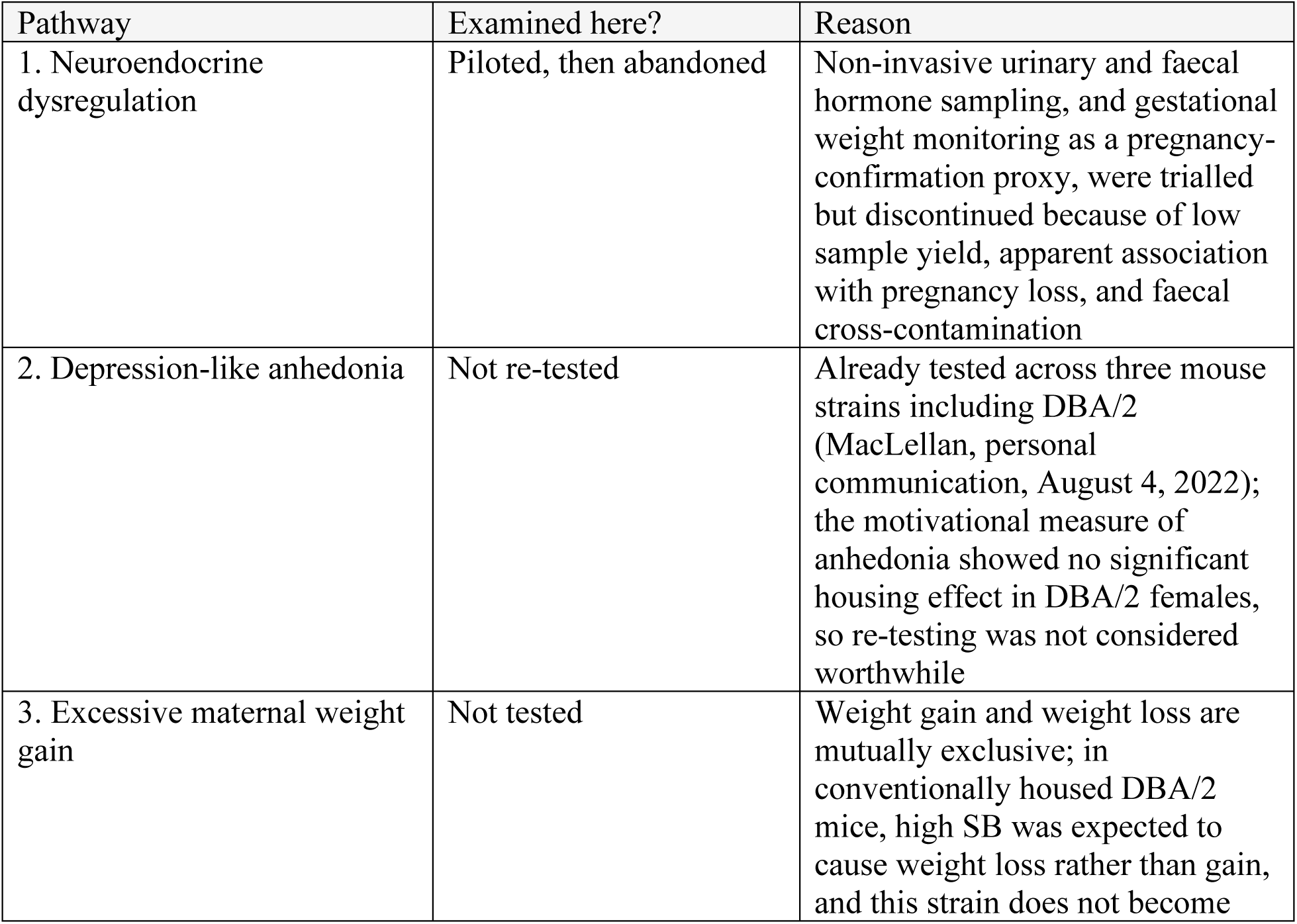
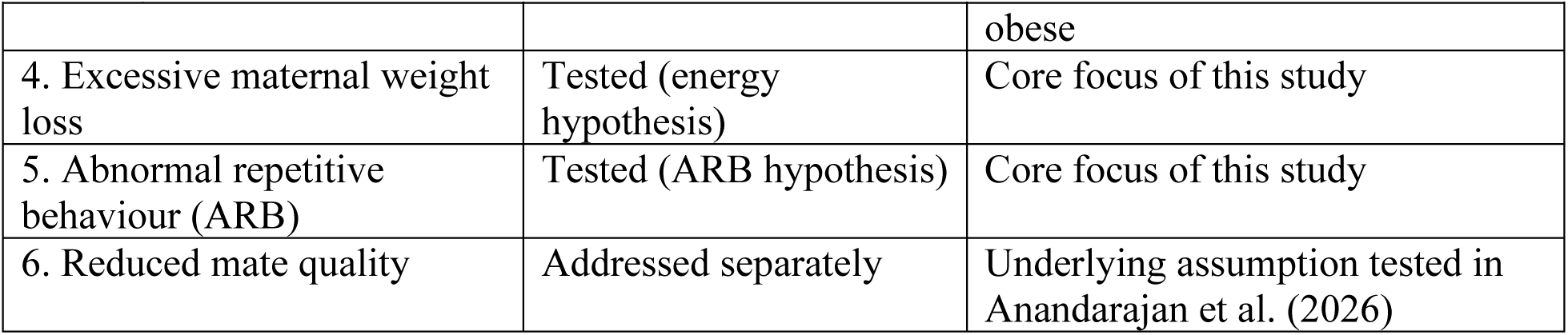
Status of the six candidate pathways linking housing quality to reproduction and maternal care.

We use "ARB" for the broad category of locomotor stereotypies, oral stereotypies and self-directed behaviours such as plucking. We use "stereotypic behaviour" (SB) for the locomotor and oral forms measured here. The two mechanisms overlap, because locomotor stereotypies can cause weight loss (Jeppesen et al., 2004; Christie et al., 2006). They are still distinct. Under the energy hypothesis, low body condition from any cause compromises reproduction. Under the ARB hypothesis, SB competes with reproduction and maternal care regardless of body condition.

### 1.1 The energy pathway

Ovulation, conception, pregnancy and lactation are among the most energetically demanding processes a female mammal undertakes (Wade & Schneider, 1992). Pre-conception body condition and gestational weight are well-established predictors of reproductive outcome (Butte & King, 2005; Rasmussen & Yaktine, 2009; Liu et al., 2016; Montvignier Monnet et al., 2022). Chronic stress can cause weight loss through increased metabolic activity (Allen et al., 2010), stress-induced anorexia (Bazhan & Zelena, 2013) and persistent SB. Being underweight carries documented reproductive costs across mammals, including infanticide in nutritionally stressed rodents and, in humans, preterm birth, low birthweight and impaired lactation.

### 1.2 The ARB/stereotypic-behaviour pathway

Barren, restrictive housing consistently predicts SB across taxa (Mason, 1991; Mason & Latham, 2004). SB is thought to reflect thwarted motivation redirected into repetitive activity (Latham & Mason, 2010). It also reflects, and may further alter, underlying central nervous system (CNS) function (Tatemoto et al., 2022). This may explain why high-SB animals can be behaviourally inflexible.

At least four non-exclusive mechanisms could link SB to poorer reproduction:

1. SB and poor reproduction share a common cause in poor welfare, such as stress or anhedonia.
2. Energetically costly SB reduces body condition (the energy pathway).
3. SB competes directly with courtship, mating or parental care for a female’s time (Mason & Burn, 2011).
4. CNS changes make high-SB females less responsive in ways that reproduction requires.

Evidence across taxa broadly supports a cost of SB to reproduction not explained by energetics alone, but it is inconsistent. SB varies in its causes and in how physically active it is. Studies also differ in design. Where high-and low-SB animals are reared identically, high-SB animals need not have had worse lives and may even breed better. Where SB is induced by housing, high-SB animals are also those with the poorer environment. Our comparison of CH and WR females is of this second, housing-manipulated kind.

### 1.3 Why this matters, and why DBA/2 mice

Many endangered mammals are bred in zoos and breeding centres where reproductive failure and abnormal behaviour limit success. Yet no study has tested, within one framework, whether SB and maternal weight loss act independently, whether one mediates the other, or whether both operate together. Even the basic claim that housing reliably affects reproduction in a fully controlled comparison is not established. Barren housing reliably induces SB in laboratory mice (Mason, 1991, 2010; Mason & Latham, 2004). DBA/2 mice are especially prone to SB and also have small litters and poor reproductive performance (Charles River

Laboratories, 1999; The Jackson Laboratory, 2024). This makes them well suited to testing links between housing, SB, body condition and reproduction.

### 1.4 Hypotheses and predictions

We test three levels of hypothesis.

1. **Housing affects reproduction.** CH females will show poorer reproductive and maternal outcomes than WR females. This must hold before mechanism is meaningful.
2. **Housing affects the candidate mechanisms.** CH females will show (a) lower BMI and (b) more SB than WR females. These are necessary, but not sufficient, conditions for either to explain a housing effect.
3. **Mechanism.** Two non-exclusive routes:

- *Energy hypothesis:* low BMI predicts smaller litters, lighter pups, poorer nests, delayed pup retrieval, less nursing, slower pup growth, earlier weaning and more infanticide. Any SB–outcome correlation should disappear once BMI is controlled for.
- *ARB hypothesis:* SB predicts longer latency to pregnancy, lower pregnancy rate, lighter pups, lower pup survival, poorer nests, delayed retrieval, less nursing and slower pup growth. These correlations should persist once BMI is controlled for.

The two hypotheses differ mainly in how each variable behaves once the other is controlled for, and we test this contrast throughout.

## 2. Methods

### 2.1 Ethics

All procedures were approved by the University of Guelph Animal Care Committee (AUP #4570) and complied with Canadian Council on Animal Care guidelines. Reporting follows ARRIVE 2.0 (Percie du Sert et al., 2020).

### 2.2 Animals and housing

We used three sequential cohorts (A, B, C) of DBA/2 mice from Charles River (Raleigh, North Carolina), obtained at 3–4 weeks of age. Each cohort had 12 pairs of sisters from different litters. Related females were housed together because they breed cooperatively, including communal nursing (König, 1993; Green et al., 2023). Pairs were randomly allocated to WR or CH, giving 6 WR and 6 CH pairs per cohort (18 of each overall). To improve low conception rates, Cohort C also received green vegetation (spinach, broccoli, lettuce), which boosts reproduction in California voles (Nelson et al., 1983; Rissman & Johnston, 1986).

Cohort A used 6 unrelated males, all in WR cages. Cohorts B and C used brother pairs from different litters, split between WR and CH by alternating assignment. Each male was housed with an adult ovariectomized CD-1 female for social contact (Gaskill & Pritchett-Corning, 2015).

CH cages contained corncob bedding, a paper cup, crinkled paper strips and cotton pads. WR cages, which reduce anxiety, depressive-like behaviour and SB in mice (Nip, 2018; MacLellan, 2023), were larger (Figure 1) and contained an igloo, running wheel, hut, tunnel, extra nesting materials, chew blocks, climbing structures, a nest box, cardboard items, hammocks and pinecones (Anandarajan et al., 2026). All items were autoclaved. Each WR cage had a detachable conventional-sized annex, joined by a tunnel, which mice were trained to enter for treats to allow low-stress capture.

**Figure 1.**
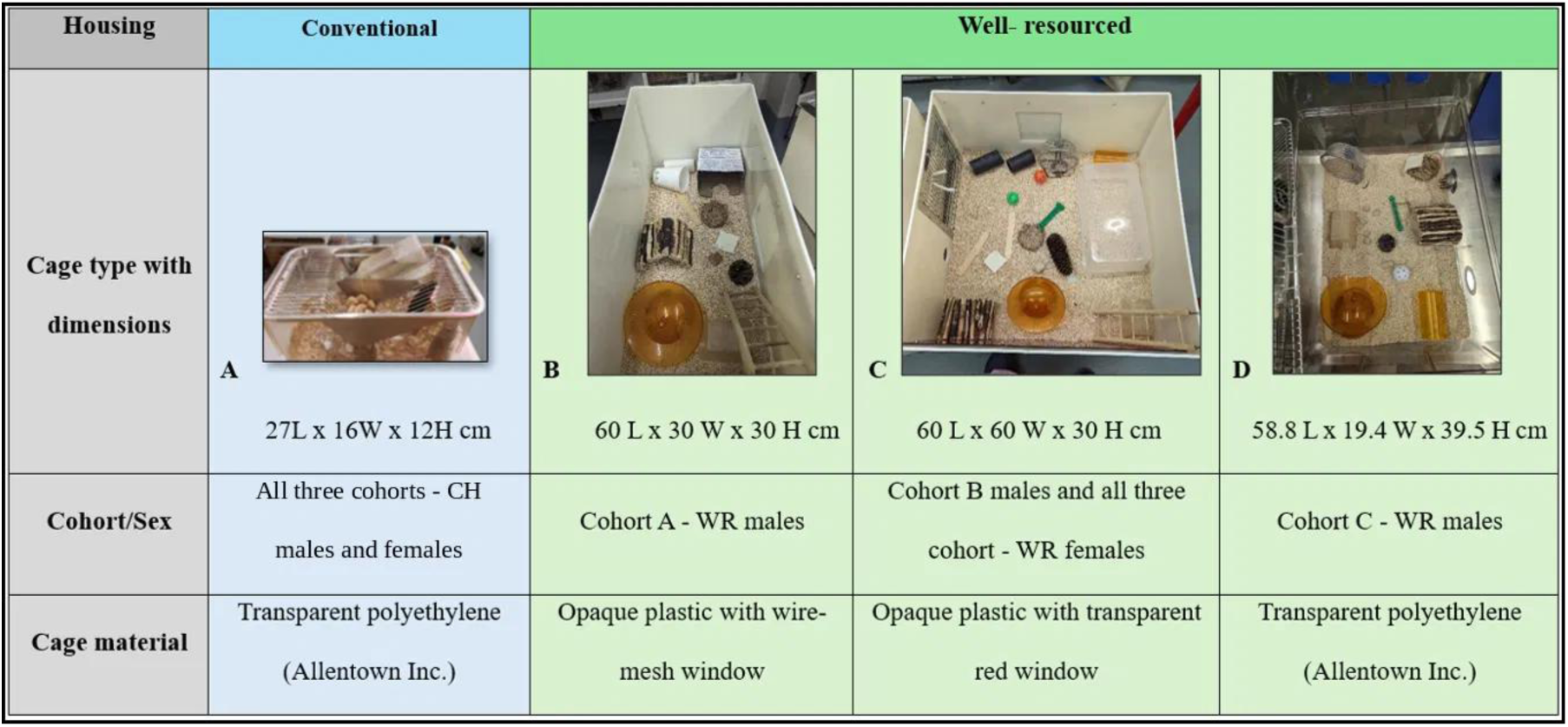
Cage types used in the study (approximately to scale). Mice were handled by cup or tunnel and habituated to 3D handling (Marcotte et al., 2021; Gouveia & Hurst, 2017). The room was kept at 21 ± 1 °C and 35–55% humidity on a reversed 14L:10D cycle (lights off 07:00). Food (Teklad Global 18% protein) and water were available *ad libitum*.

### 2.3 Behavioural observation

Home-cage time budgets were collected at 10–11 weeks of age, before mating, over four consecutive days. Scans began two hours into the dark phase and ran for four hours, at least five minutes apart (cf. Tilly et al., 2010). Two observers recorded the first behaviour seen for each mouse, using a predefined ethogram (Table 2). Maternal care and SB were scored by one observer from PND 17 to PND 28, 09:00–13:00 (cf. Bechard & Mason, 2010). Earlier maternal care could not be scored because dams stayed out of sight in the nest. Because WR mice had more places to hide, all time budgets were calculated as proportions of visible scans. SB was scored in the same sessions as maternal care, so we could test whether time in SB came at the expense of care.

**Table 2.**
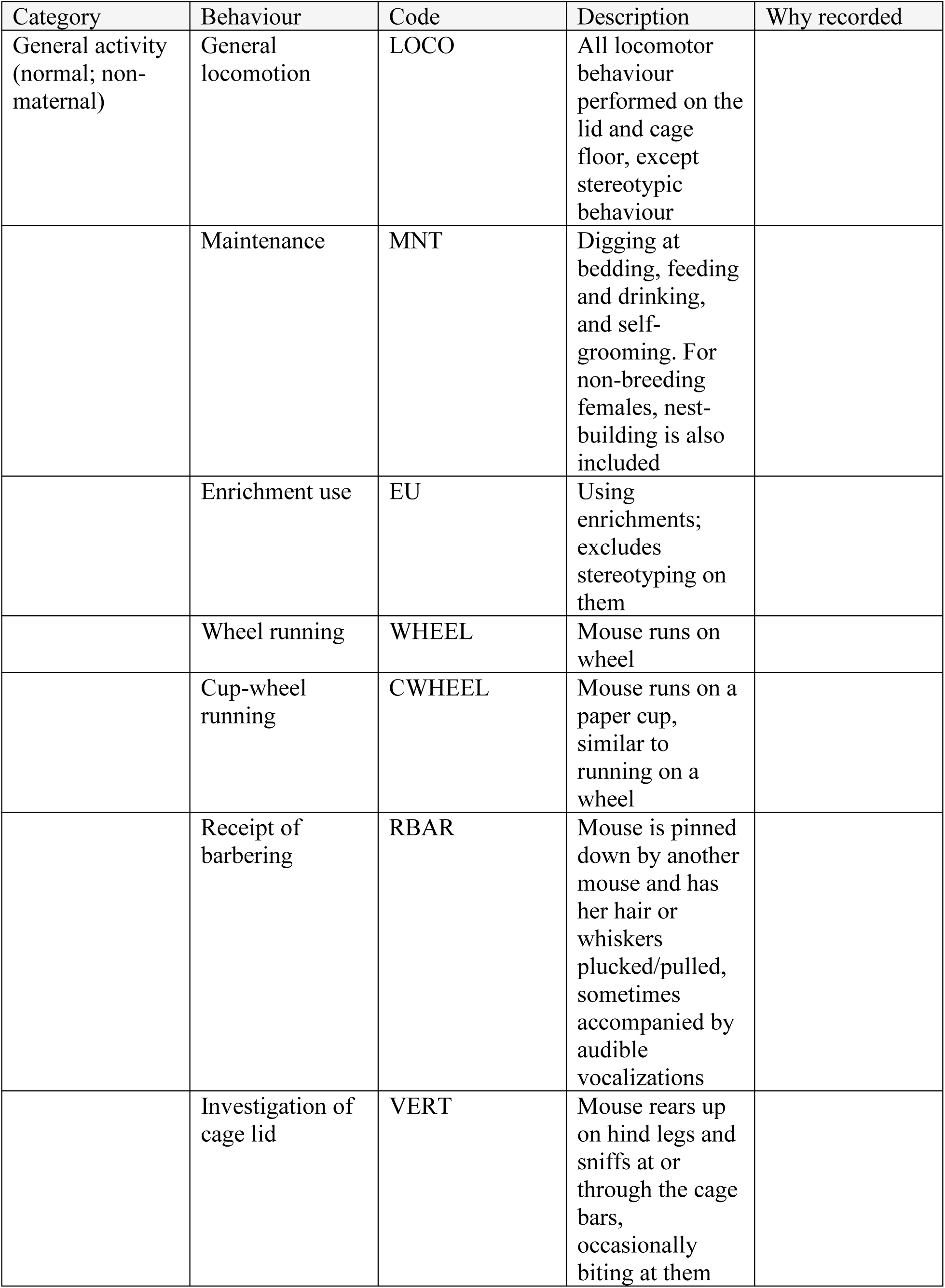
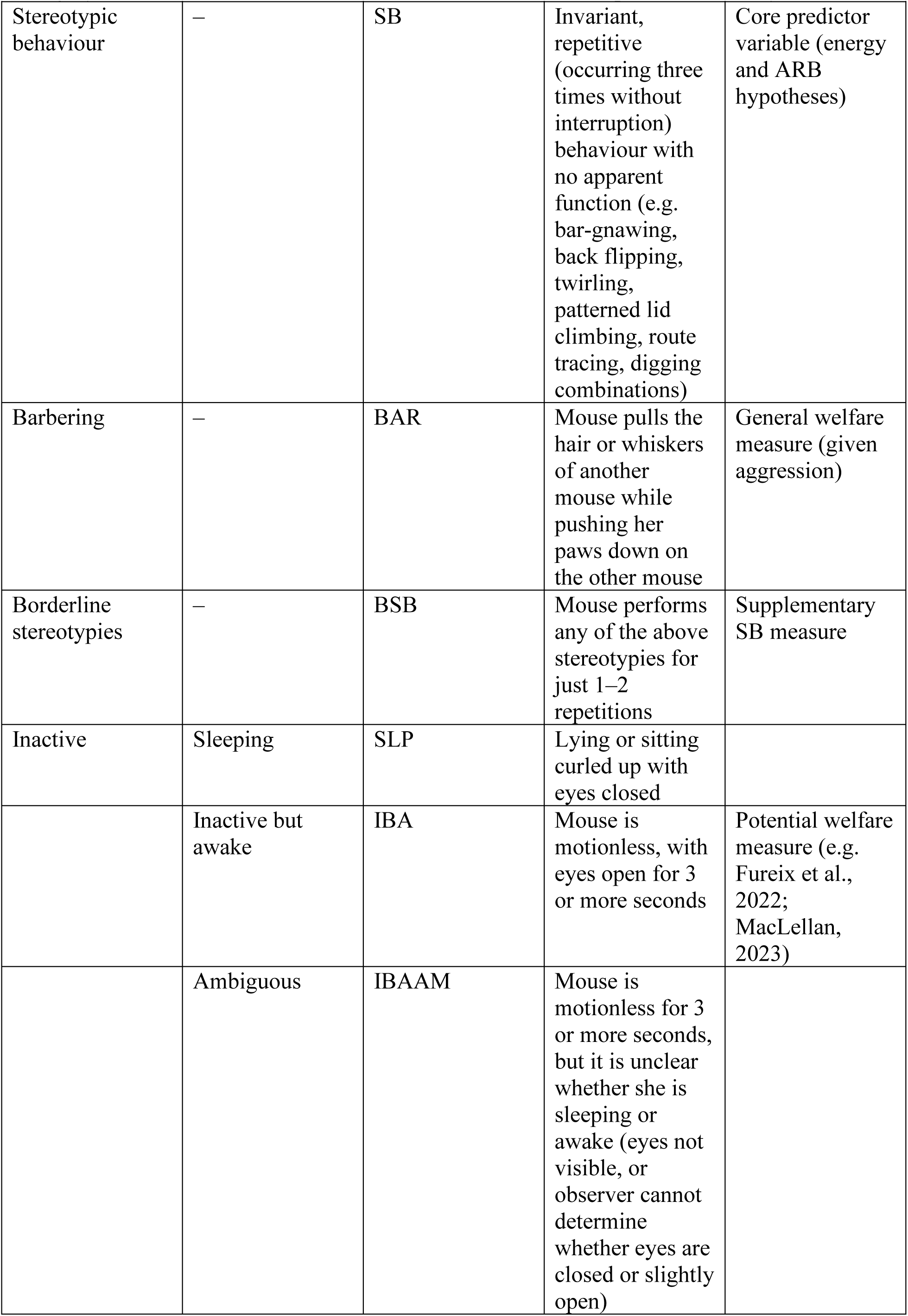
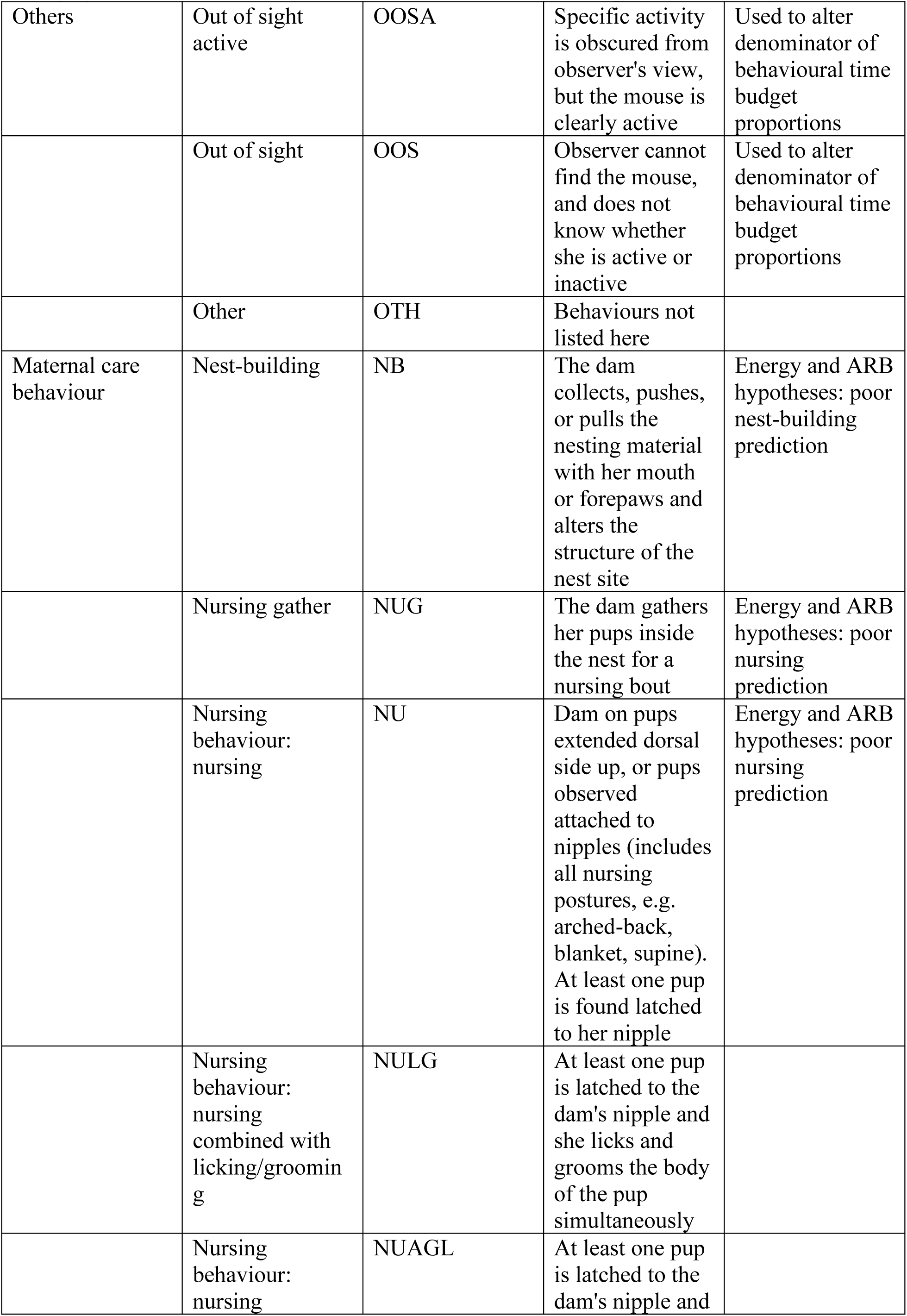
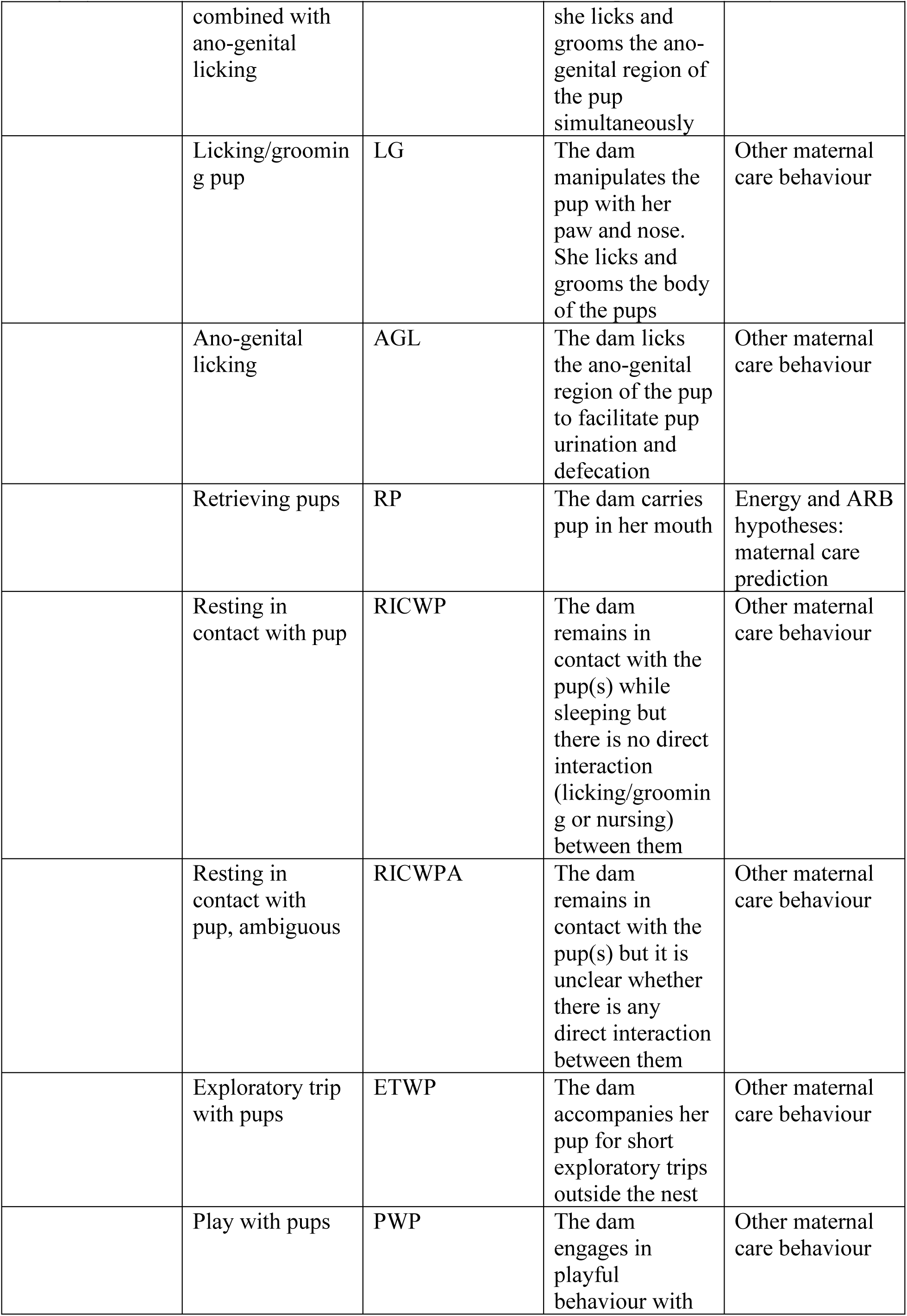
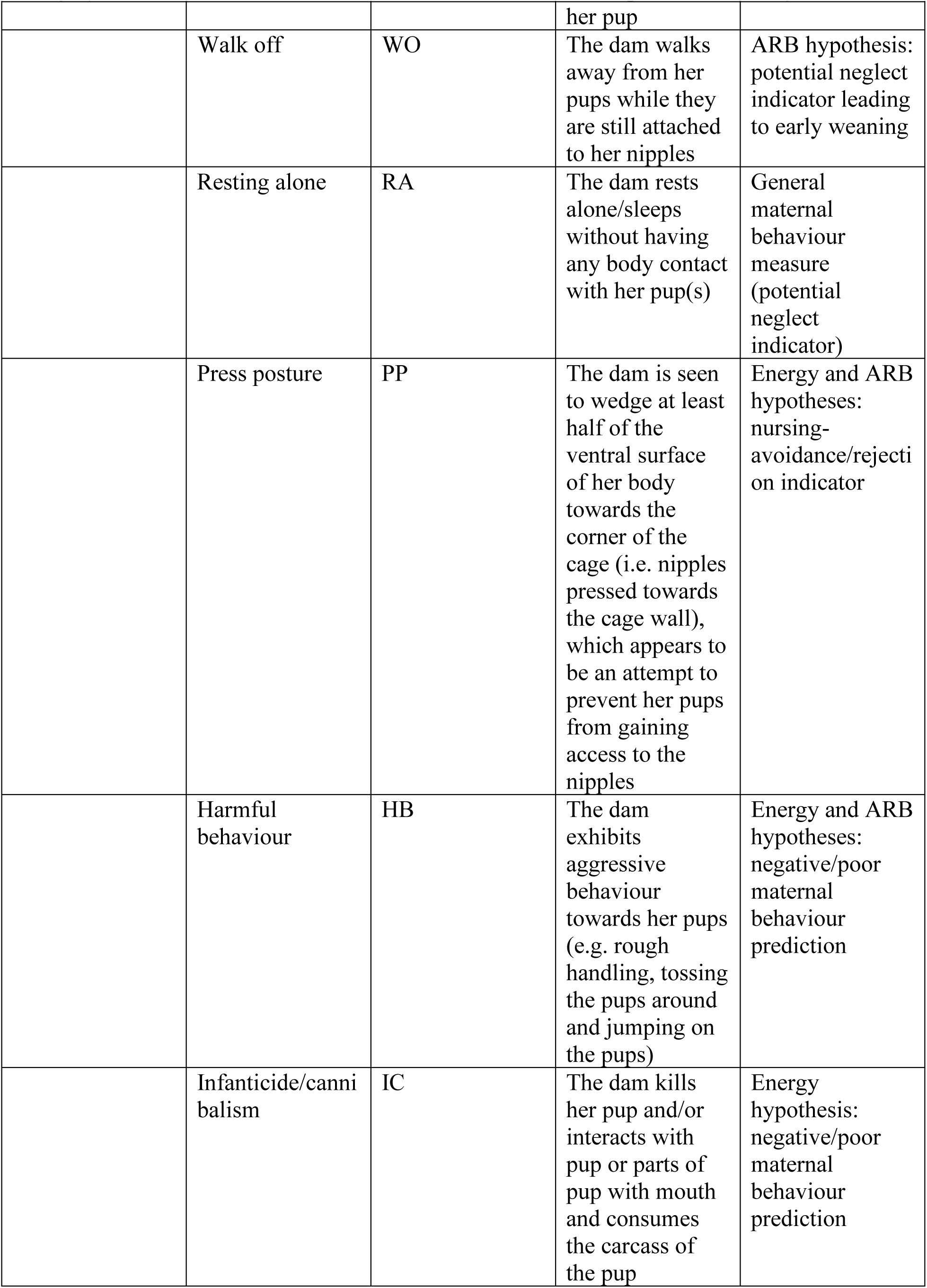
Ethogram used for home-cage and maternal care observations in female mice (Latham & Mason, 2004; Shoji & Kato, 2006; Bechard & Mason, 2010; Chourbaji et al., 2011; Gaskill & Pritchett-Corning, 2015; Weber, 2015; Nip, 2018; Orso et al., 2019; Preguiça, 2017).

### 2.4 Body mass index

BMI (weight / length²) reflects body fat more accurately than weight alone (MacLellan, 2023). Females were weighed the day before mating and on PND 16, at peak lactational demand (Knight et al., 1986; McDonald & Nielsen, 2006). Nose-to-anus length was measured from ventral photographs in ImageJ (version 1.52a, National Institutes of Health, USA) by an observer blind to housing (Figure 2).

**Figure 2.**
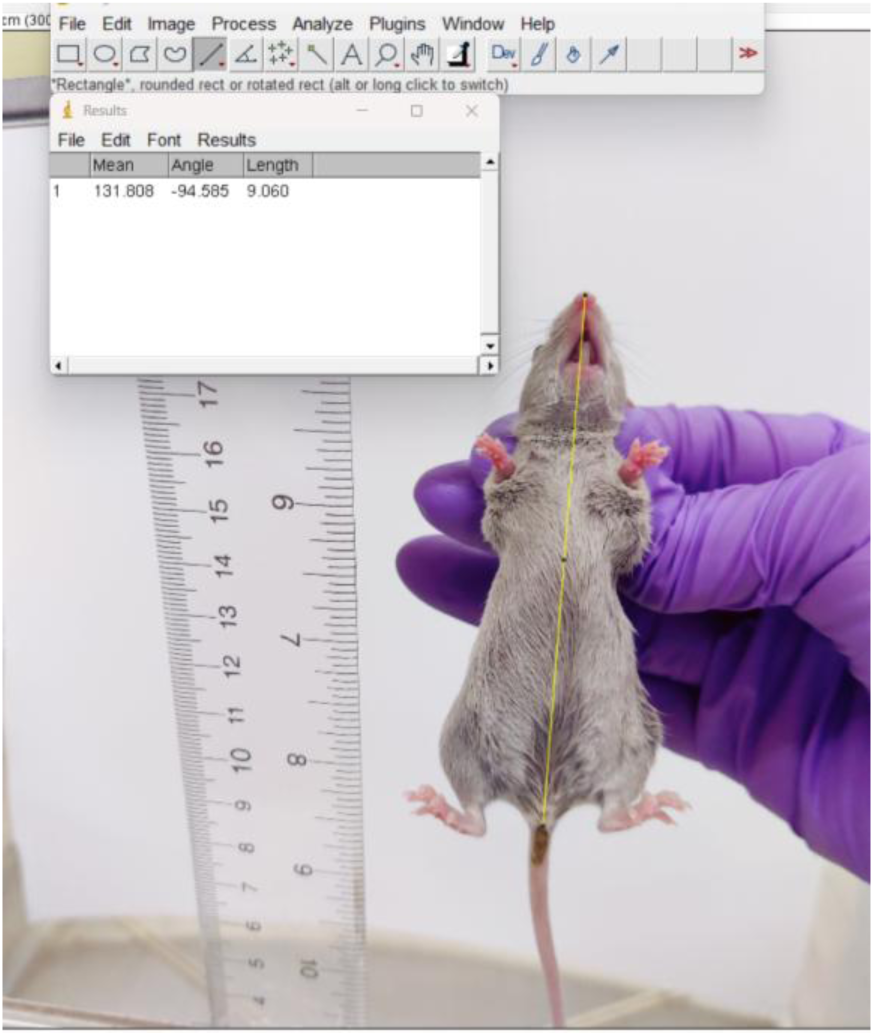
Body length (nose-to-anus) measured in ImageJ from ventral photographs.

### 2.5 Oestrus induction and mating

Each sister pair was exposed to soiled bedding from WR and CH males for three days, to induce oestrus via the Whitten effect (Whitten, 1956) and familiarize females with male odours. A UV urine detector was used to select bedding with similar urine content from both cage types.

In Cohort A, one oestrous female per pair took a sexual incentive motivation test and was then paired with a male for four hours. This test did not predict mating outcomes and added substantial handling, so it was dropped. Because birth rates were low, Cohorts B and C used continuous trio-housing instead. Each sister pair was housed with a similarly reared male for 21 days, starting 72 hours after oestrus induction (Cohort B) or after mate-choice tests at that same time point (Cohort C; Anandarajan et al., 2026). Trios were watched for two hours after introduction. Mating was recorded only when the full sequence of courtship, intromission and ejaculation was seen. Plugs were not checked, to avoid stressing females.

Pregnancy was assessed from abdominal enlargement, starting ten days after first observed mating (cf. Pang et al., 2014). Latency to pregnancy was the time from trio-housing to unambiguous pregnancy. Females not pregnant within 35 days were excluded from pregnancy-latency and pup analyses but kept in mating-rate analyses.

Non-invasive urinary and faecal hormone sampling and gestational weight monitoring were piloted in early cohorts but abandoned. Sample yields were low, sampling appeared to be associated with pregnancy loss, and faecal samples were cross-contaminated.

### 2.6 Maternal and offspring measures Nest score

Nests were photographed from above and from the side on PND 7 and scored 0–5 (0 = untouched material, 5 = complete dome) by a blinded observer (Gaskill, Karas, et al., 2013; Figures 3 and 4).

**Figure 3.**
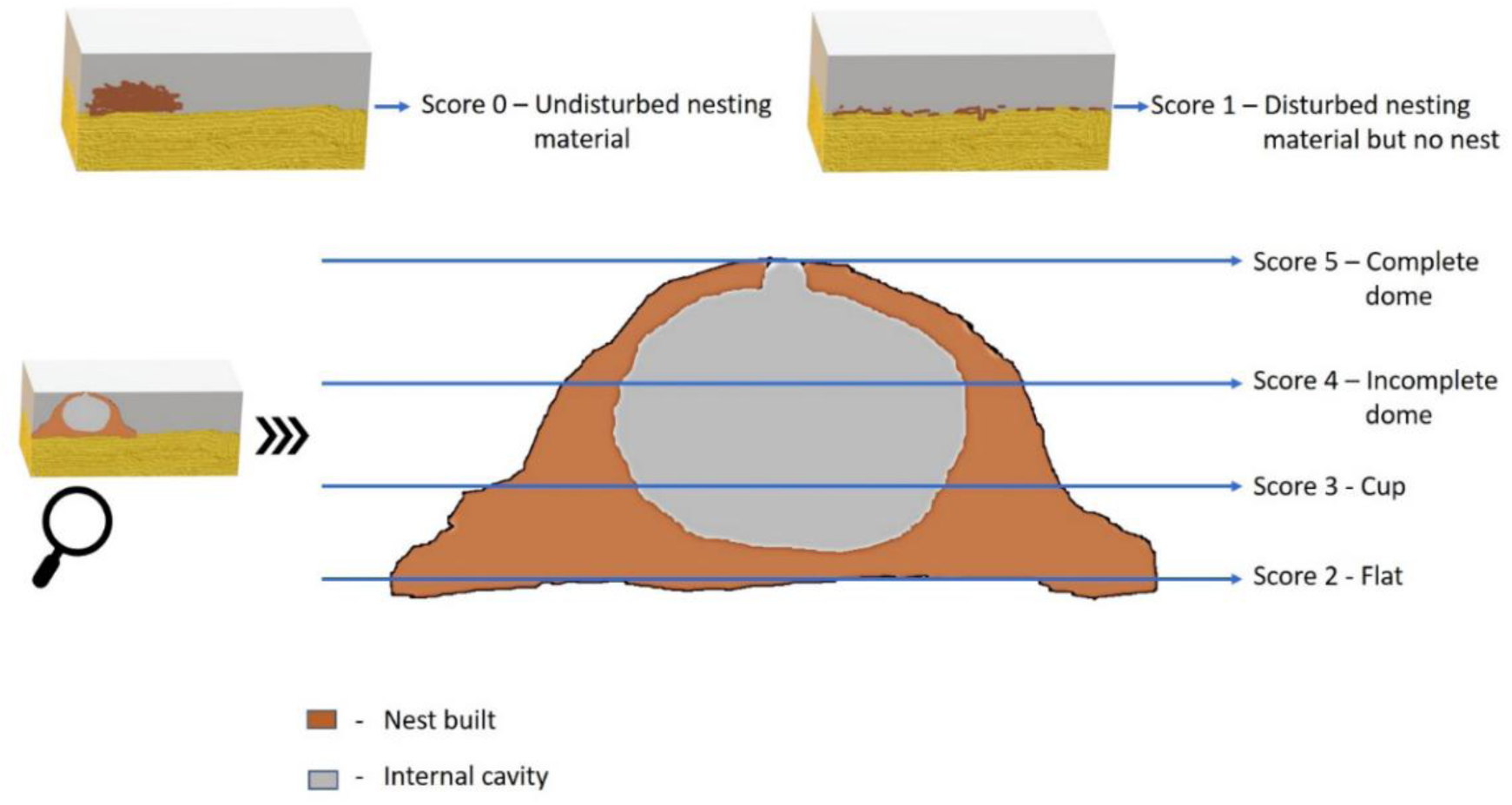
Nest scoring scale: material undisturbed (0), disturbed but no nest built (1), flat nest (2), cup-shaped nest (3), incomplete dome (4) and complete dome (5) (Gaskill, Karas, et al., 2013).

**Figure 4.**
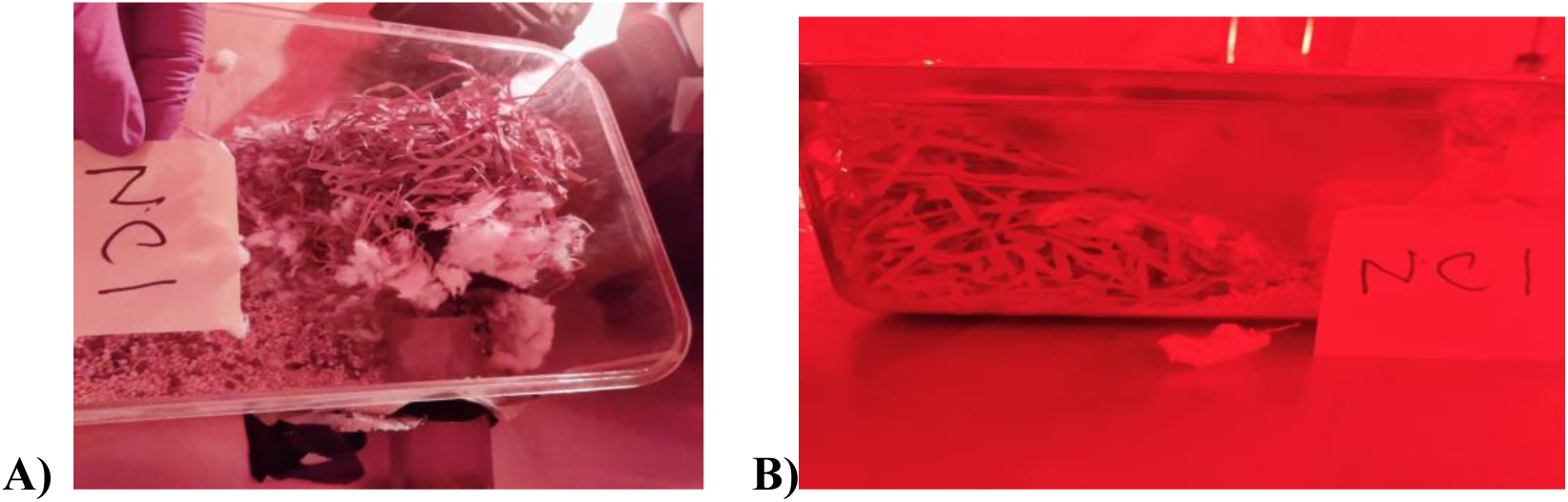
Nest photographs taken on PND 7 for scoring: (A) top view and (B) side view. The nest shown scores 5. Other measures were mating rate, latency to pregnancy, pregnancy rate, pups born, pups weaned and pup survival.

#### Pup retrieval

On PND 7, two pups were placed 15 cm from the nest in the opposite corner of the home cage, behind a barrier (modified from Meek et al., 2001). After the barrier was removed, retrieval was scored live for 15 minutes by a blinded observer. Pups not retrieved were assigned 900 s. WR dams were tested in their annex, where they mostly nested.

#### Nursing and weaning

Nursing frequency came from the PND 17–28 scans. The last day nursing was seen indexed weaning timing (Curley et al., 2009), alongside walk-off frequency.

#### Pup growth

Pups were weighed to 0.01 g on PND 0, 16, 21 and 28, and litter means were used.

#### Milk yield

Weigh-suckle-weigh tests on PND 16 were excluded because nursing after separation was inconsistent.

## 3 Statistical analysis

Analyses were run in R 4.1.3 (R Core Team, 2022), with α = 0.05. They addressed three questions in turn: does housing affect reproductive and maternal outcomes; does housing affect BMI and SB; and do BMI or SB account for any housing effect. Cohort was a fixed effect in every model. Table 3 lists the model used for each outcome.

**Table 3.**
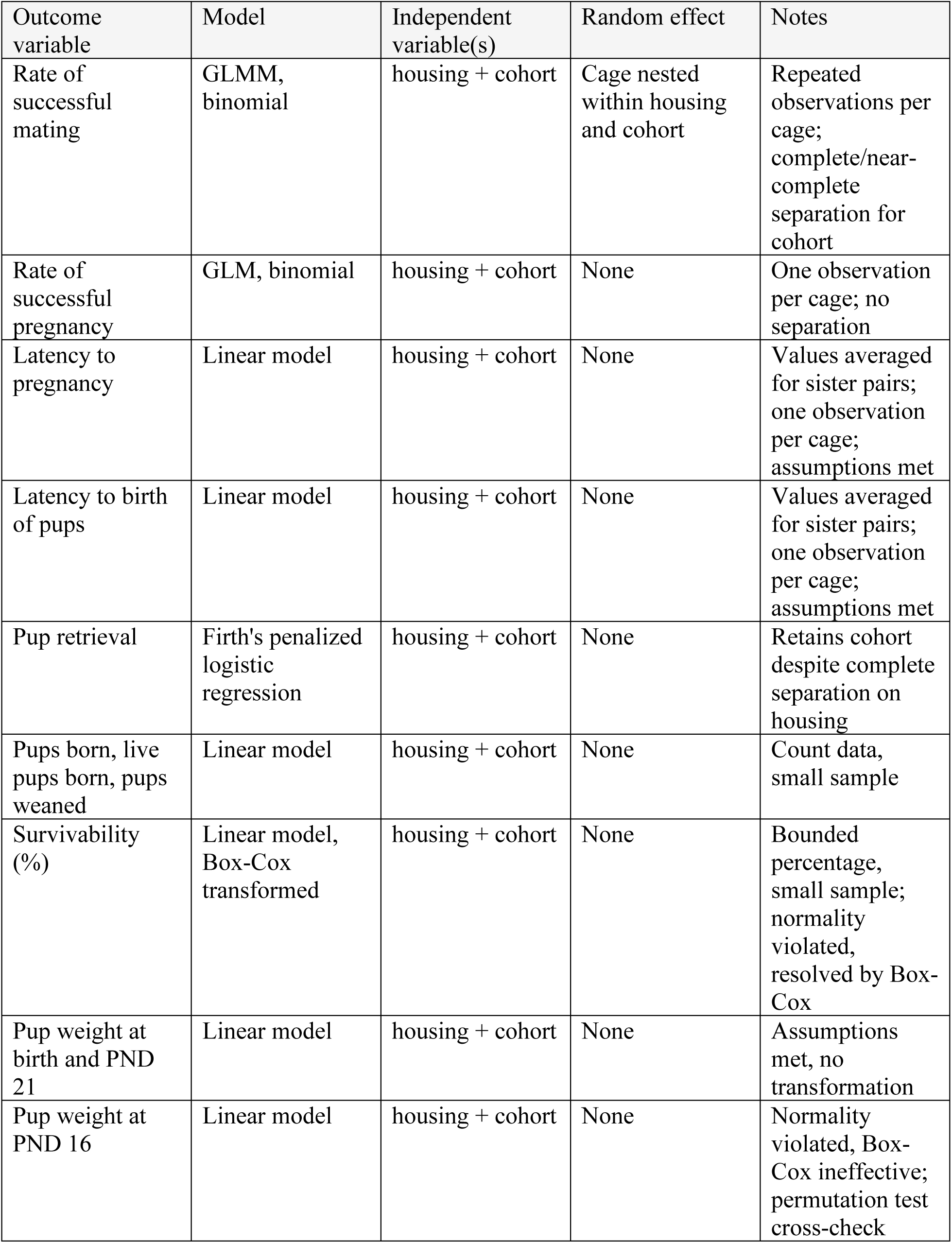
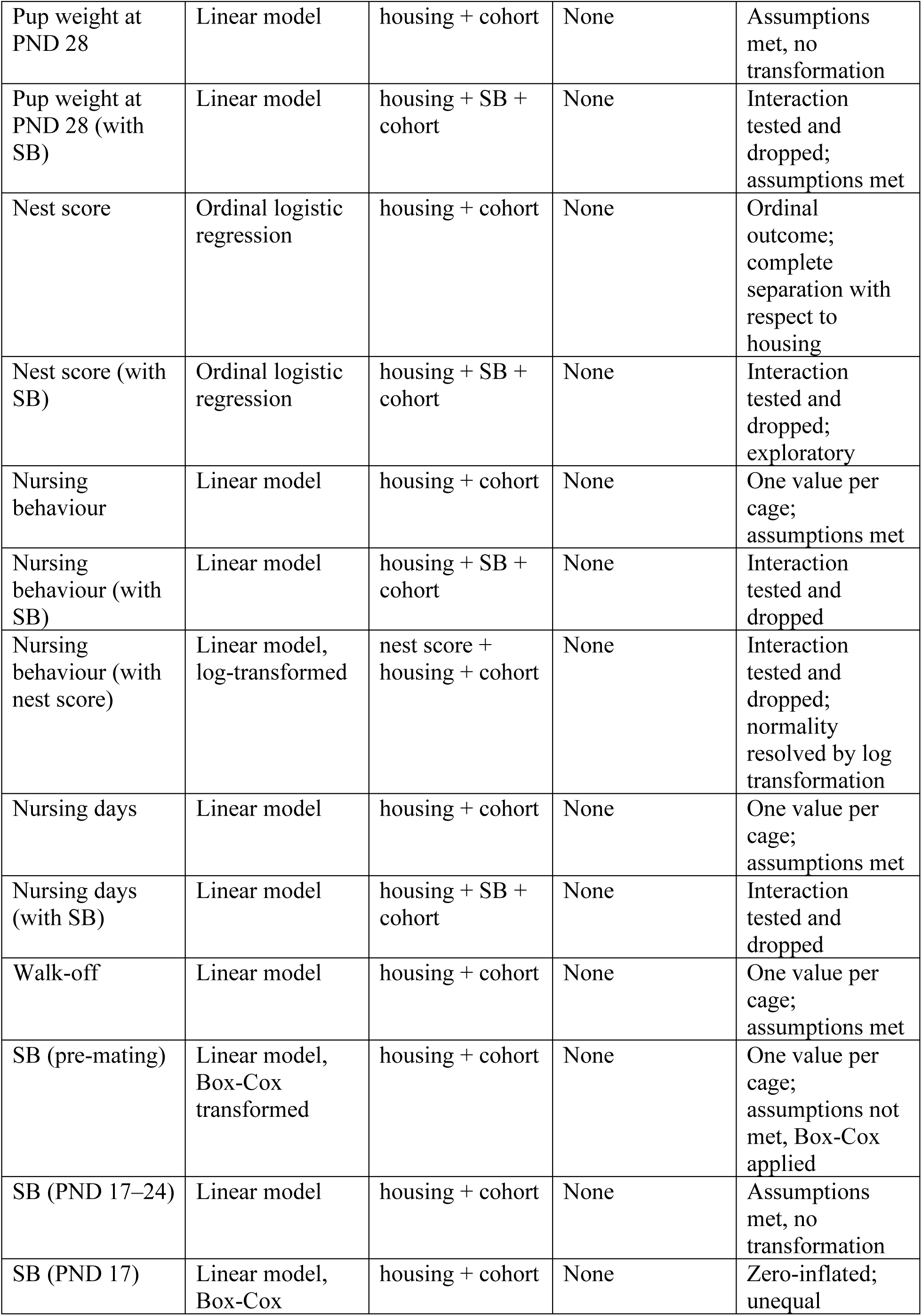
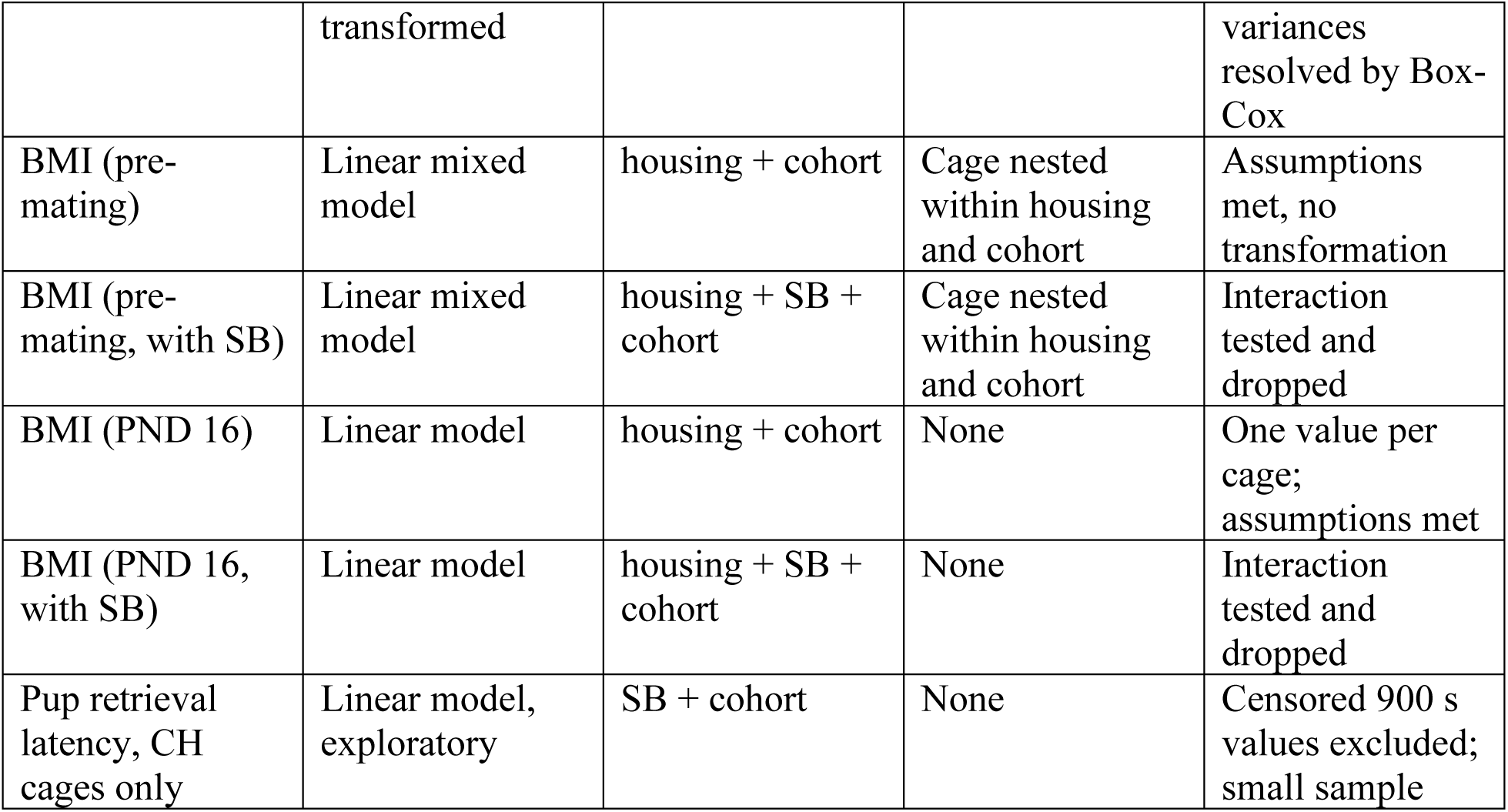
Statistical models used to test the effect of housing on reproductive, maternal care and behavioural outcomes.

Repeated measures within cages used linear mixed models with cage nested within housing and cohort. Outcomes with one value per cage used linear models. Normality and variance were checked with Shapiro–Wilk and Levene’s tests. Non-normal outcomes were Box-Cox or log transformed. Where transformation failed, the untransformed model was kept and checked with a permutation test. Binary outcomes used binomial GLMMs or GLMs. Where housing perfectly predicted the outcome, Firth’s penalized logistic regression was used. Nest score was analyzed by ordinal logistic regression.

For mechanism, each affected outcome was modelled with housing, cohort and SB and/or BMI. A housing × covariate interaction was tested first and dropped if non-significant. We then asked whether the housing effect weakened once the covariate was added. As an exploratory check, pup retrieval latency was modelled against SB within CH cages only (n = 7), the only cages with retrievals. SB and maternal care were analysed for PND 17–24, because all pups were naturally weaned by PND 24.

## 4 Results

### 4.1 Housing effects on reproduction and maternal care

Of the 36 breeding pairs/trios established (18 CH, 18 WR; 6 CH and 6 WR per Cohort), 17 gave birth to live pups (7 CH, 10 WR), varying by Cohort as shown in **Table 4**. In two Cohort C litters (both WR), the single pup born was found dead on postnatal day 1 however, so no surviving pups were available from those cages; all subsequent nest score, maternal behaviour, and pup weight analyses are therefore based on the remaining 15 cages with surviving litters (7 CH, 8 WR).

**Table 4:** Breeding pairs/trios and cohort-specific procedural differences.

| Cohort | Male housing | Oestrus induction / pairing | Mating protocol | Other procedures running alongside | Cages with pups born (CH / WR) | Cages with surviving pups (CH / WR) | Mean pups / cage | Median | Range | Litters of $\leq 2$ pups |
| --- | --- | --- | --- | --- | --- | --- | --- | --- | --- | --- |
| **A** | 6 unrelated males, all WR (not split by housing) | Vaginal lavage + cytology to confirm oestrus | Single-day pairing — only one female per sister pair (whichever came into oestrus first) was paired with the male until 5pm, then male removed | Urinary hormone sampling (daily isolation up to 2h, 5 days + GD13) and GD10+ weight monitoring; sexual incentive motivation test, after which males were allowed with females for mating | CH 1/6, WR 2/6 | CH 1/6, WR 2/6 | 3.0 | 2.0 | 2–5 | 2 of 3 |
| **B** | Brother pairs, differentially housed WR/CH | Bedding-exposure (Whitten effect), 3 days pre-familiarization | Continuous trio-housing with male for 21 days | Faecal sampling piloted (glitter-marked); milk yield test separation extended to 3h | CH 4/6, WR 2/6 | CH 4/6, WR 2/6 | 6.5 | 6.0 | 3–13 | 0 of 6 |
| C | Brother pairs,<br>differentially<br>housed WR/CH | Same<br>bedding-exposure<br>protocol as B | Continuous trio-housing, male<br>selected from prior<br>mate-choice test | Green vegetation added<br>(spinach/broccoli/lettuce);<br>faecal sampling continued;<br>milk yield test moved to<br>PND14 | CH 2/6,<br>WR 6/6 | CH<br>2/6,<br>WR<br>4/6 | 5.1 | 5.5 | 1–<br>9 | 2<br>of<br>8 |

Number of pups born seem to improve after Cohort A, on two fronts: the proportion of pairs/trios having pups increased (3 of 12 in Cohort A, versus 6 of 12 in Cohort B and 8 of 12 in Cohort C — roughly 25% → 50% → 67%), and of those who did have pups, mean pups per cage also increased (3.0 → 6.5 → 5.1). This coincided with several changes at once — dropping the stressful urinary/weight sampling, moving to continuous trio-housing with both sisters rather than mating only one female per cage as in Cohort A, and adding green vegetation in Cohort C -- so it is not possible to say which specific change (or combination) drove the improvement -- future researchers to test individually.

#### 4.1.1 Reproductive success and litter outcomes

Housing quality did not significantly affect any of the following reproductive outcomes (see **Table 5**): rate of successful mating (Cohort: χ²₂ = 12.23, p = 0.002), latency to become pregnant (Cohort: F₂,₂₇ = 5.35, p = 0.011), rate of successful pregnancy (Cohort: χ²₂ = 1.33, p = 0.514), latency to give birth (Cohort: F₂,₁₃ = 1.98, p = 0.178), number of pups born (Cohort: F₂,₁₃ = 0.81, p = 0.468), number of live pups born (Cohort: F₂,₁₃ = 0.70, p = 0.514), number of pups weaned (Cohort: F₂, ₁₃ = 1.34, p = 0.295), survivability of pups (Cohort: F₂, ₁₃ = 3.19, p = 0.075), average pup birth weight (Cohort: F₂, ₁₁ = 1.98, p = 0.184), average pup weight at PND 16 (Cohort: F₂, ₁₁ = 0.16, p = 0.855), and average pup weight at PND 21 (Cohort: F₂, ₁₁ = 0.06, p = 0.941).

However, at PND 28, though pup weight did not differ significantly by housing, a trend effect was noticed, with WR pups tending to weigh more than CH pups (see **Table 5**) (Cohort: F₂, ₁₁ = 0.04, p = 0.962).

Analyses of nest scores revealed a significant difference between WR and CH females, with WR females having higher scores (housing: χ²₁ = 9.92, p = 0.002; Cohort: χ²₂ = 1.87, p = 0.393) (**Figure 3.5**)

**Figure 5:**
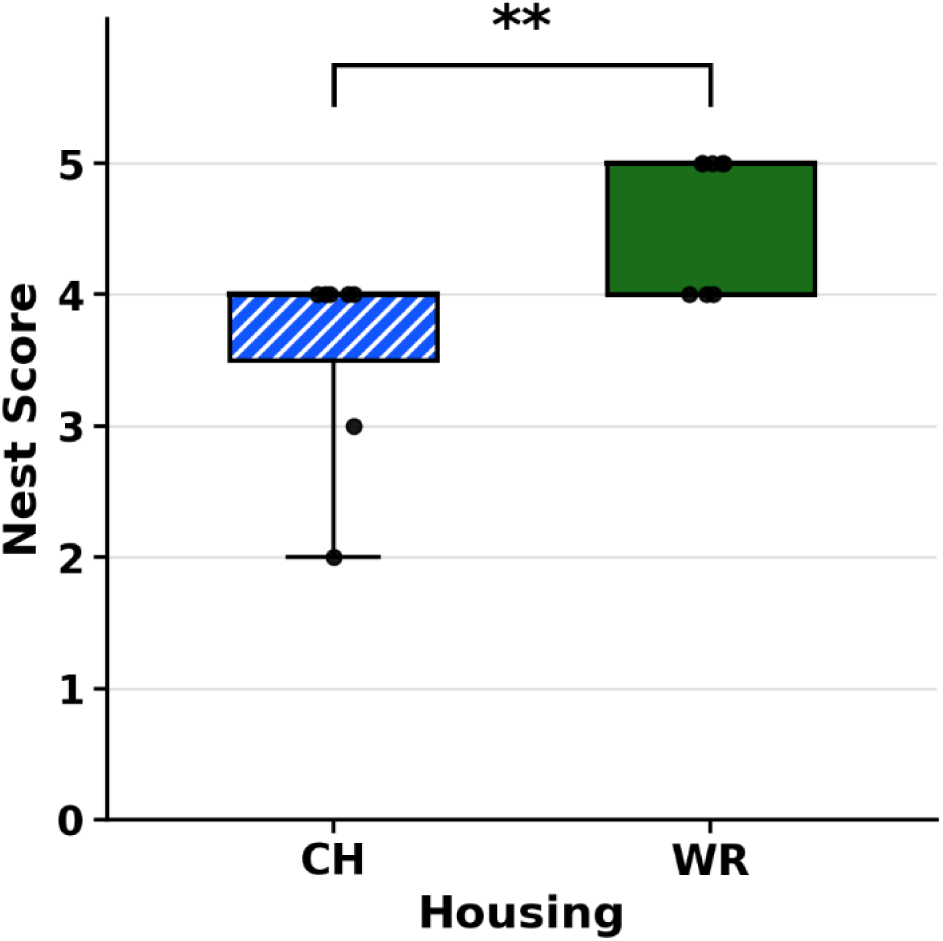
Nest score by housing quality. Medians and interquartile ranges are presented, with individual cage values overlaid. ** = p ≤ 0.01.

Pup retrieval success also differed significantly between housing conditions (χ²_1_ = 12.29, p < 0.001), but unexpectedly with 100% success in CH females (7/7) and 0% success in WR females (0/8) (Cohort: χ²_2_ = 0.07, p = 0.965) (**Figure 6**)

**Figure 6.**
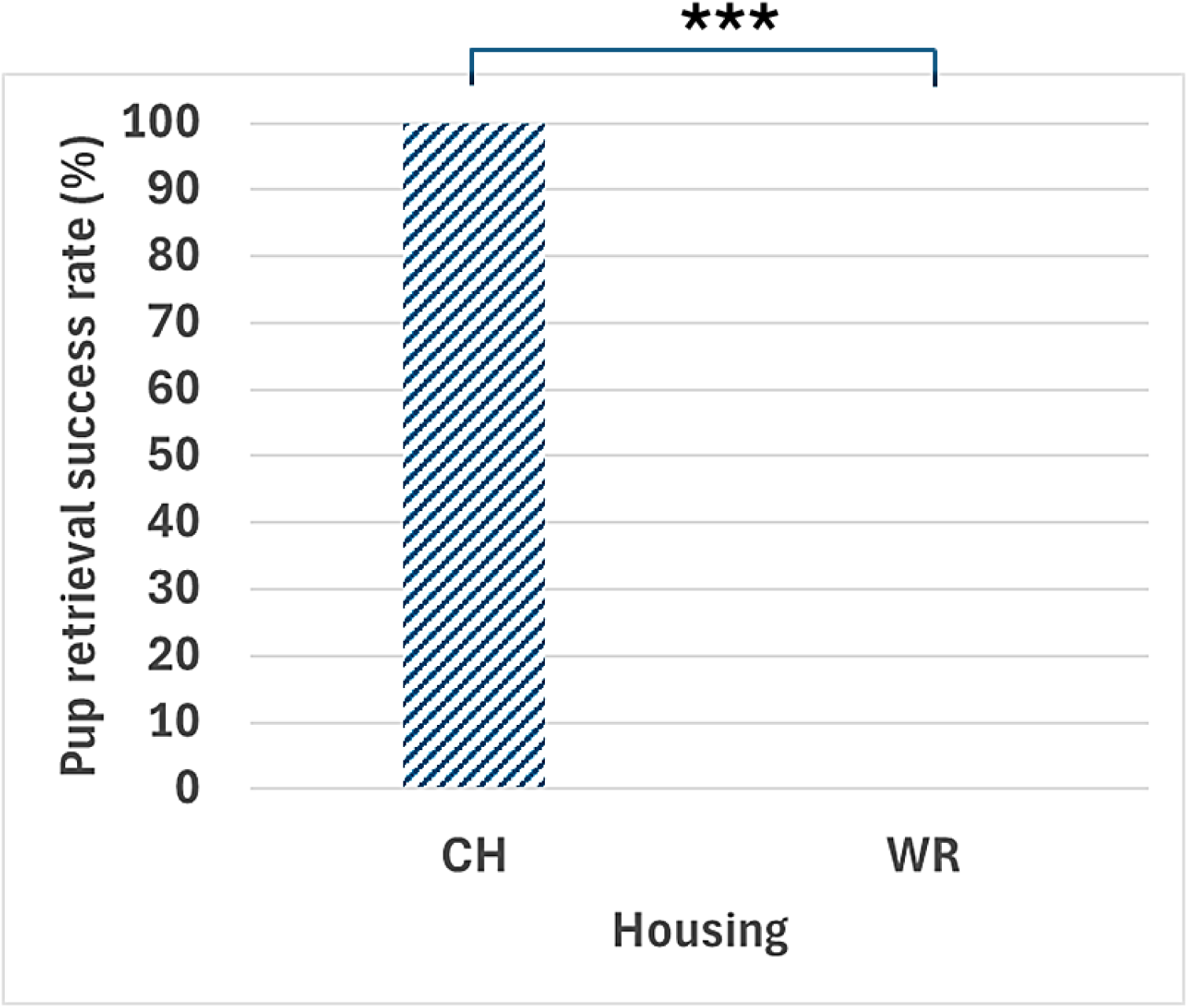
Pup retrieval success rate for CH and WR housing. Pup retrieval rates are given in percentage. **\*\*\*** = p ≤ 0.001.

**Table 5:**
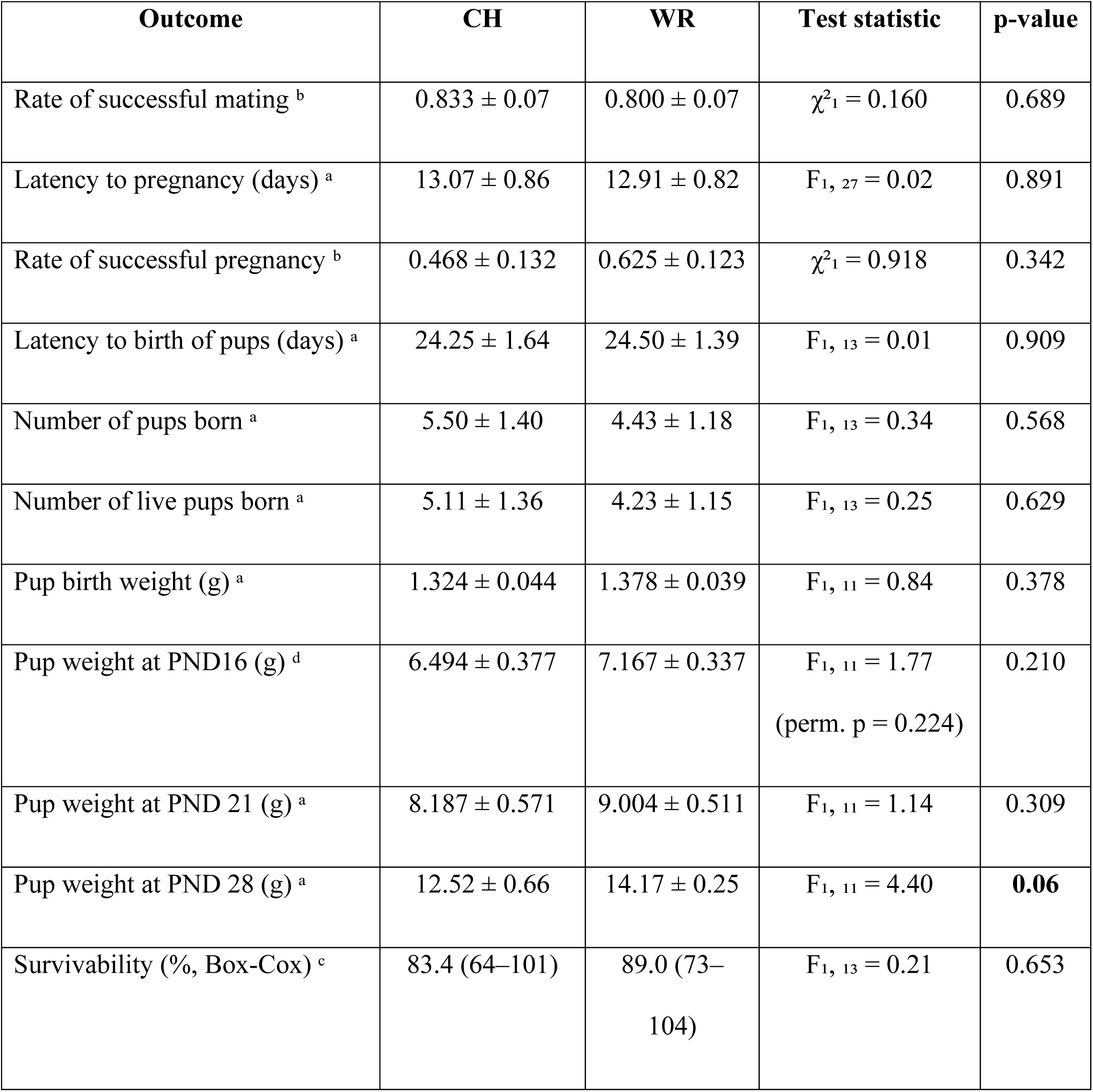
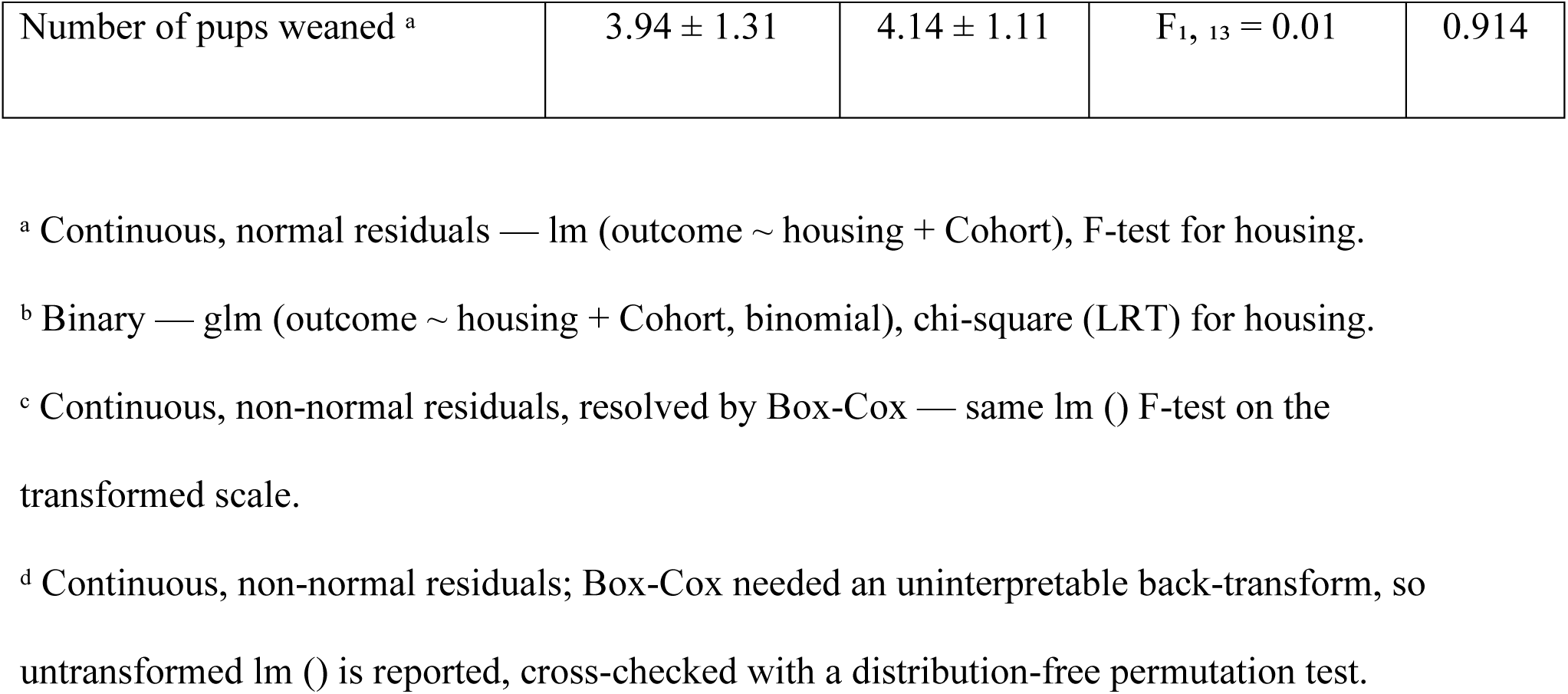
Reproductive parameters for WR and CH females.

| Outcome | CH | WR | Test statistic | p-value |
| --- | --- | --- | --- | --- |
| Rate of successful mating <sup>b</sup> | 0.833 ± 0.07 | 0.800 ± 0.07 | $\chi^2_1 = 0.160$ | 0.689 |
| Latency to pregnancy (days) <sup>a</sup> | 13.07 ± 0.86 | 12.91 ± 0.82 | $F_{1, 27} = 0.02$ | 0.891 |
| Rate of successful pregnancy <sup>b</sup> | 0.468 ± 0.132 | 0.625 ± 0.123 | $\chi^2_1 = 0.918$ | 0.342 |
| Latency to birth of pups (days) <sup>a</sup> | 24.25 ± 1.64 | 24.50 ± 1.39 | $F_{1, 13} = 0.01$ | 0.909 |
| Number of pups born <sup>a</sup> | 5.50 ± 1.40 | 4.43 ± 1.18 | $F_{1, 13} = 0.34$ | 0.568 |
| Number of live pups born <sup>a</sup> | 5.11 ± 1.36 | 4.23 ± 1.15 | $F_{1, 13} = 0.25$ | 0.629 |
| Pup birth weight (g) <sup>a</sup> | 1.324 ± 0.044 | 1.378 ± 0.039 | $F_{1, 11} = 0.84$ | 0.378 |
| Pup weight at PND16 (g) <sup>d</sup> | 6.494 ± 0.377 | 7.167 ± 0.337 | $F_{1, 11} = 1.77$<br>(perm. p = 0.224) | 0.210 |
| Pup weight at PND 21 (g) <sup>a</sup> | 8.187 ± 0.571 | 9.004 ± 0.511 | $F_{1, 11} = 1.14$ | 0.309 |
| Pup weight at PND 28 (g) <sup>a</sup> | 12.52 ± 0.66 | 14.17 ± 0.25 | $F_{1, 11} = 4.40$ | **0.06** |
| Survivability (%; Box-Cox) <sup>c</sup> | 83.4 (64–101) | 89.0 (73–104) | $F_{1, 13} = 0.21$ | 0.653 |
| Number of pups weaned <sup>a</sup> | 3.94 ± 1.31 | 4.14 ± 1.11 | F<sub>1, 13</sub> = 0.01 | 0.914 |

<sup>a</sup> Continuous, normal residuals — lm (outcome ~ housing + Cohort), F-test for housing.
<sup>b</sup> Binary — glm (outcome ~ housing + Cohort, binomial), chi-square (LRT) for housing.
<sup>c</sup> Continuous, non-normal residuals, resolved by Box-Cox — same lm () F-test on the transformed scale.
<sup>d</sup> Continuous, non-normal residuals; Box-Cox needed an uninterpretable back-transform, so untransformed lm () is reported, cross-checked with a distribution-free permutation test.

#### 4.1.2 Effects of housing quality on nursing behaviour, nursing days and ‘walk-off’ behaviour

##### Nursing behaviour

Analyses of nursing behaviour revealed a significant effect of housing (F_1, 11_ = 7.80, p = 0.018), with WR females showing a significantly greater proportion of visible observations spent nursing over the whole lactation period than CH females (Cohort: F_2, 11_ = 0.28, p = 0.761) (**Figure 3.7**)

**Figure 7:**
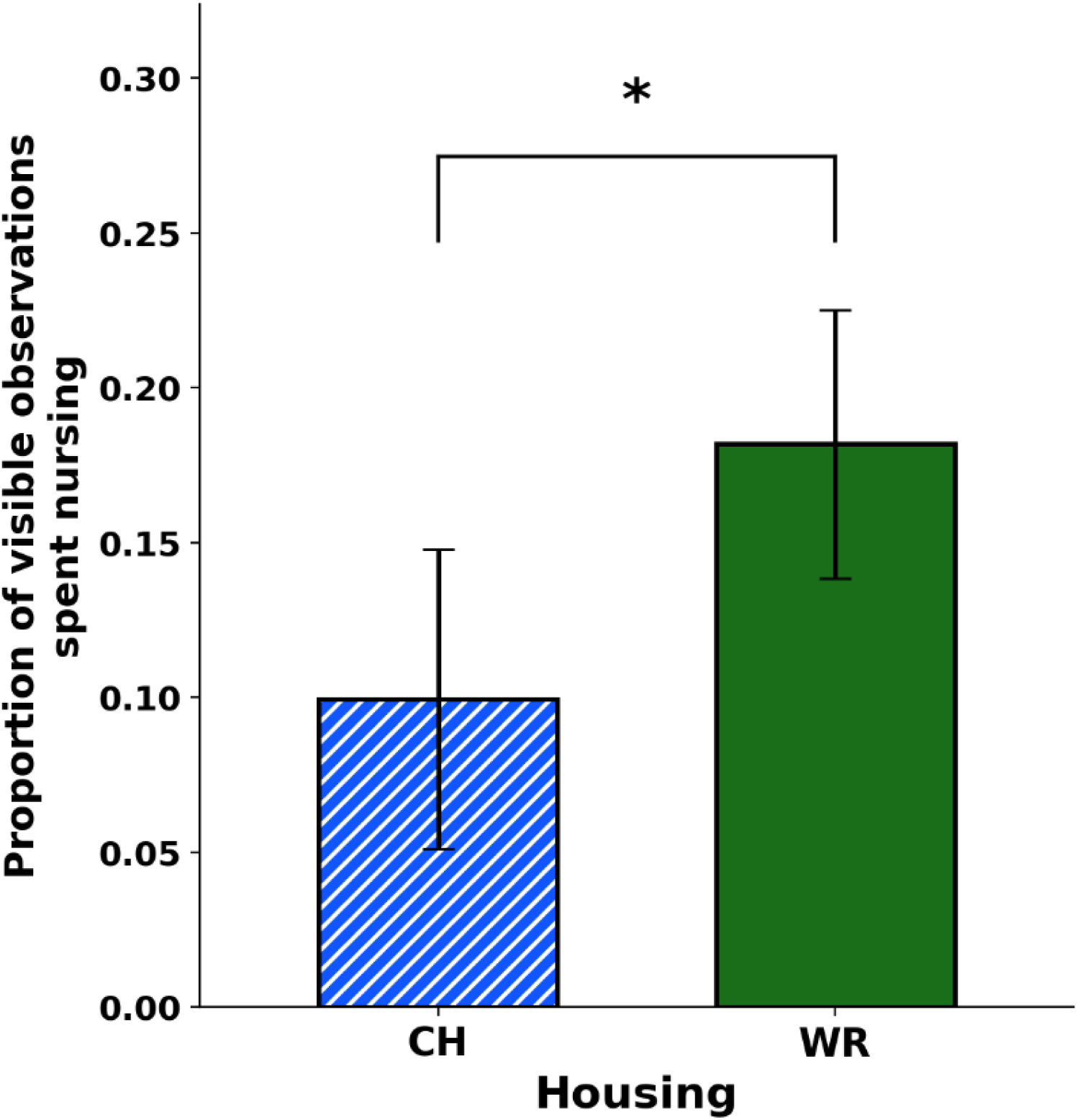
Proportion of visible observations spent nursing by the females based on housing conditions. Mean ± standard errors are presented. * = p ≤ 0.05.

##### Nursing days to assess early weaning

Housing significantly predicted the number of nursing days, an indicator of early weaning, with WR females nursing for more days than CH females (F₁, ₁₁ = 6.98, p = 0.023) (Cohort: F₂, ₁₁ = 2.47, p = 0.130), suggesting earlier weaning of pups by CH dams (**Figure 8**).

**Figure 8:**
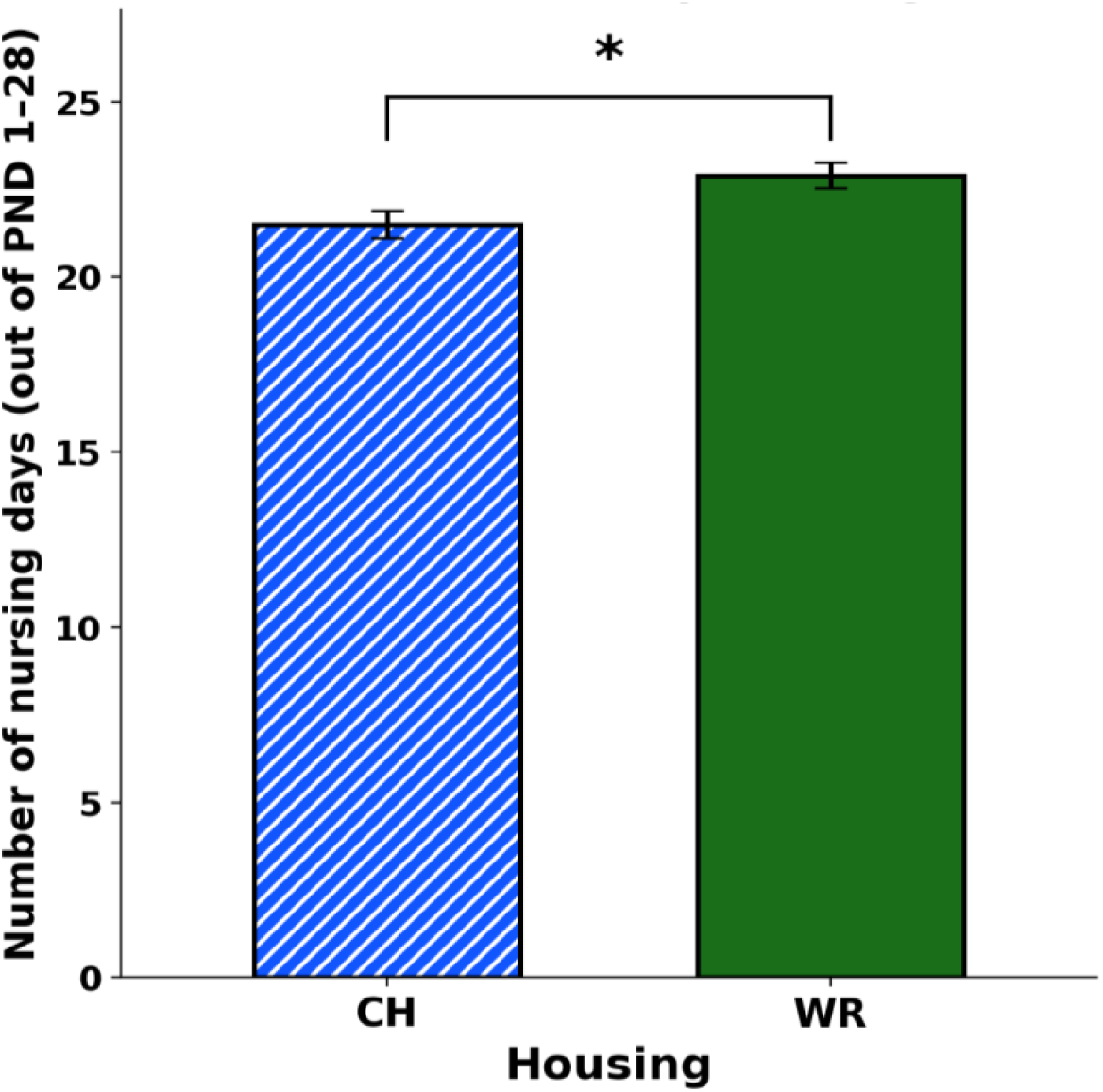
Number of nursing days (out of PND 1 to 28) by the females based on housing conditions. Mean ± standard errors are presented. * = p ≤ 0.05.

##### ’Walk-off’ behaviour to assess natural weaning

Housing had no significant effect on walk-off behaviour, with CH and WR females showing near-identical proportions (CH = 0.0029 ± 0.001 and WR = 0.0028 ± 0.001) of visible observations spent walking off from pups while pups were still sucking milk (F₁, ₁₁ = 0.68, p = 0.427) (Cohort: F₂, ₁₁ = 14.95, p = 0.0007)

##### Nursing behaviour and nest score

To test whether nursing behaviour covaried with nest score, a linear model was fit with nest score, housing, and Cohort as predictors of nursing (log-transformed to meet assumptions) A housing × nest score interaction was tested first and was not significant (F₁, ₉=0.0055, p=0.942), so it was dropped and the additive model is reported. In the additive model, nest score showed a trend towards predicting nursing (F₁, ₁₀ = 3.40, p = 0.095), with higher nest scores associated with more nursing. Nursing rose from 0.091± 0.019 at a nest score of 3 to 0.168 ± 0.029 at a nest score of 5 — nearly a two-fold increase.

### 4.2 Stereotypic behaviour and BMI: Potential effects of housing quality that could impair reproduction

#### 4.2.1 Stereotypic behaviour

Before the mating period, analyses of the proportion of visible observations spent exhibiting SB by females revealed a significant effect of housing quality (F_1, 32_ = 82.62, p < 0.001), with CH females spending significantly more time (almost 9.6 times more) stereotyping than WR females. (Cohort: F_2, 32_ = 7.12, p = 0.003) (**Figure 9**).

**Figure 9:**
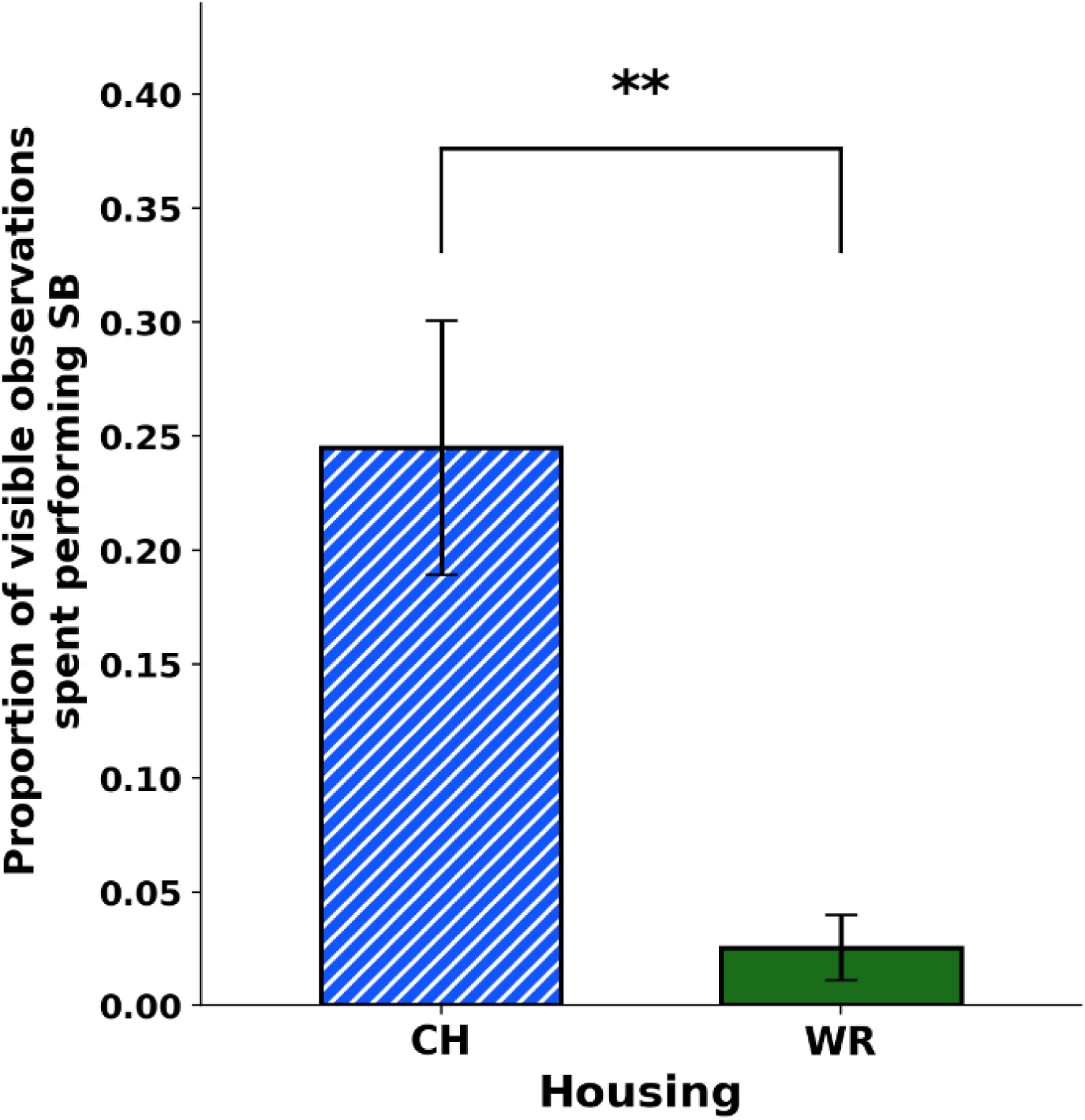
Proportion of visible observations spent in stereotypic behaviour by the females before the mating period. Mean ± standard errors are presented. ** = p ≤ 0.01.

On PND 17, analyses of the proportion of visible observations spent exhibiting SB by lactating females also revealed a significant effect of housing quality (F₁, _1_₁ = 14.10, p = 0.003), with CH females again exhibiting more SBs than WR females, who now rarely performed SB at all (**Figure 10**).

**Figure 10:**
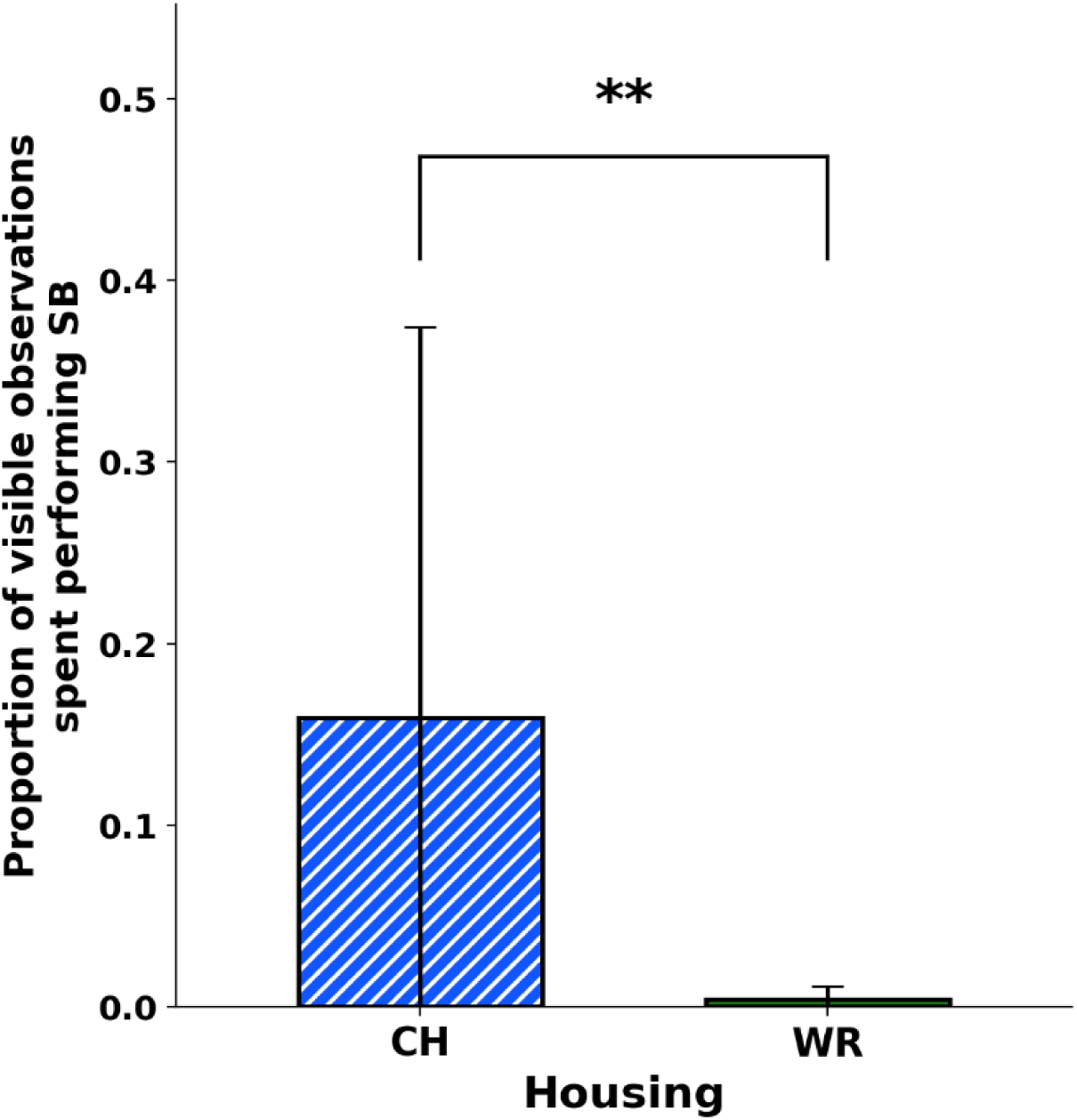
Proportion of stereotypic behaviour exhibited by the females on PND17 based on housing conditions. Mean ± standard errors are presented. ** = p ≤ 0.01.

During later lactation (PND 17 to PND 24), housing still significantly predicted overall stereotypic behaviour (F_1, 11_ = 16.04, p = 0.002), with CH females showing higher levels (5.1 times more) than WR females (Cohort: F_2, 11_ = 6.64, p = 0.013) **(Figure 11).**

**Figure 11:**
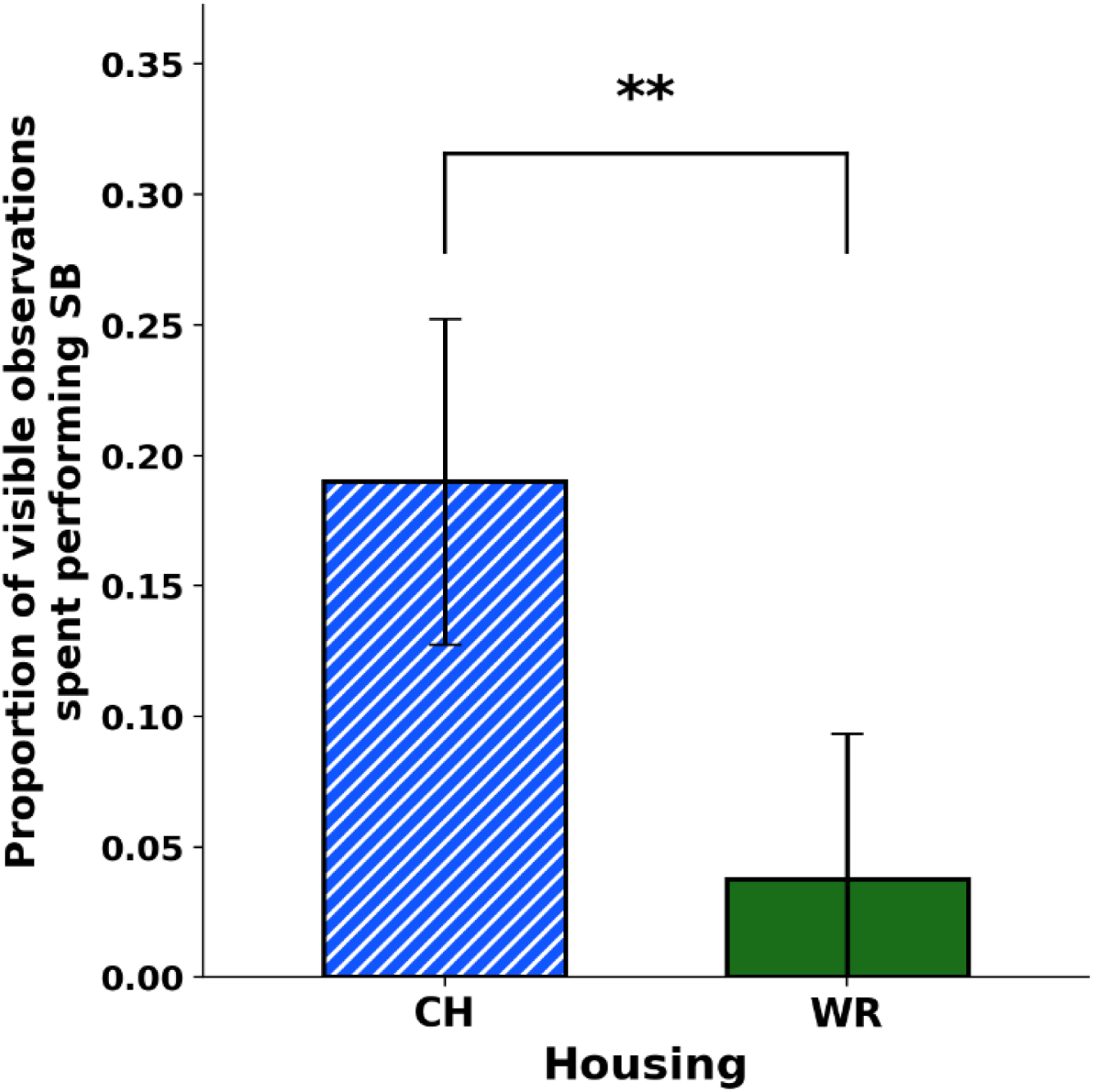
Proportion of stereotypic behaviour exhibited by the females from PND17 to PND 24 based on housing conditions. Mean ± standard errors are presented. ** = p ≤ 0.01.

#### 4.2.2 BMI

Housing quality had no significant effect on BMI, neither in the periods before mating (Cohort: F₂, ₃₂ = 60.26, p < 0.001) nor on post-natal day 16 (PND 16) (Cohort: F₂, ₁₁ = 7.19, p = 0.010) (**Table 6**)

**Table 6:** BMI of WR and CH females.

| Outcome | CH | WR | Test Statistic | p-value |
| --- | --- | --- | --- | --- |
| Pre-mating BMI <sup>a</sup> | 0.257±0.037 | 0.266±0.036 | $F_{1, 32} = 2.88$ | 0.100 |
| BMI at PND 16 of lactating females <sup>a</sup> | 0.324±0.06 | 0.308±0.048 | $F_{1,11} = 0.01$ | 0.923 |

<sup>a</sup> Continuous, normally distributed data — F-test (linear model).

To test whether, despite this, SB predicted BMI pre-mating, a linear mixed model was used with housing and SB and their interaction plus Cohort as fixed effect and with cage as a random effect nested within housing. The housing X SB interaction was not significant (F_1, 30_ = 0.14, p = 0.709), so it was dropped from the model. In the resulting additive model, neither housing (F_1, 31_ = 1.18, p = 0.285) nor SB (F_1, 31_ = 0.02, p = 0.897) significantly predicted BMI (Cohort: F_2, 31_ = 53.67, p < 0.001).

Likewise, at PND 16, the interaction (housing × SB) was not significant (F_1, 9_ = 1.02, p = 0.339), so it was dropped. In the resulting additive model, neither housing (F_1, 10_ = 0.06, p = 0.807) nor SB at PND 17 (F_1, 10_ = 0.12, p = 0.738) significantly predicted BMI at PND16 (Cohort: F_2, 10_ = 6.36, p = 0.017).

Overall, housing thus did not significantly affect BMI, indicating that BMI cannot serve as a mediator of any housing effect on reproductive and maternal care outcomes.

### 4.3 Does SB predict reproduction and maternal care?

Overall, the following aspects of reproduction were thus compromised by CH conditions: nest quality, time spent nursing, nursing days and (a trend) pup weight at PND 28; but pup retrieval was unexpectedly more diligent by CH mothers than WR. The relationships between SB and nest quality, time spent nursing, pup weight at PND 28 and (exploratively) pup retrieval times were therefore analyzed to assess whether SB mediated the housing effects.

#### Nest score

To test whether SB predicted nest score, an ordinal logistic regression was fitted with nest score as the outcome and housing, stereotypic behaviour (SB), and Cohort as predictors. A housing × SB interaction term was tested first and was not significant (χ²_1_ = 0.03, p = 0.869), so it was dropped and the additive model reported below. Housing (χ²_1_ = 1.72, p = 0.189) now did not significantly predict nest score (the significant effect of housing observed when modelled alone - - χ²_1_ = 9.92, p = 0.002 -- being attenuated once SB was included), while SB showed a trend towards significance (χ²_1_ = 2.94, p = 0.086), with higher SB associated with lower nest scores (Cohort: χ²_2_ = 3.12, p = 0.211). This analysis should be interpreted with caution, however, as two of the four nest score categories contained only a single observation each, limiting the stability of the parameter estimates.

#### Pup weight at PND 28

To test whether stereotypic behaviour predicted pup weight at PND 28, a linear model was fitted with housing, SB, and Cohort as predictors. A housing × SB interaction term was tested first and was not significant (F_1, 9_ = 0.02, p = 0.879), so it was dropped from the reported model. In the additive model, SB did not significantly predict pup weight at PND 28 (F_1, 10_ = 0.05, p = 0.823) but now neither did housing now that SB was in the model (F_1, 10_ = 2.12, p = 0.176) (Cohort: F_2, 10_ = 0.01, p = 0.987).

#### Nursing behaviour

To test whether SB predicted nursing behaviour, a linear model was fitted with housing, SB, and Cohort as predictors. A housing × SB interaction term was tested first and was not significant (F _1, 9_ = 0.06, p = 0.813), so it was dropped from the reported model. In the additive model, SB did not significantly predict nursing behaviour (F_(1,10)_ = 0.90, p = 0.366). However, the significant effect of housing observed when modelled alone (F_1, 11_ = 7.80, p = 0.018) was no longer significant once SB was included (F_1, 10_ = 1.09, p = 0.321) (Cohort: F_2, 10_ = 0.37, p = 0.698).

#### Nursing days to assess early weaning

To test whether SB predicted nursing days, a linear model was fitted with housing, SB, and Cohort as predictors. A housing × SB interaction term was tested first and was not significant (F₁, ₉ = 2.43, p = 0.154), so it was dropped and the additive model is reported. In the additive model, SB again did not significantly predict nursing days (F₁, ₁₀ = 2.47, p = 0.147). Notably, again here, the significant effect of housing observed when modelled alone (F₁, ₁₁ = 6.98, p = 0.023) was no longer significant once SB was included (F₁, ₁₀ = 0.34, p = 0.573) (Cohort: F₂, ₁₀ = 1.32, p = 0.309).

#### Pup retrieval time

As an exploratory, *post hoc* check on whether SB predicted the promptness of pup retrieval among CH cages, linear models were fitted with SB and Cohort as predictors for two retrieval measures: latency to retrieve first pup and latency to bring all pups to warmer area. SB did not significantly predict latency to retrieve the first pup (F_1, 3_ = 0.24, p = 0.657) nor latency to bring pups to the warm area (F_1, 3_ = 0.14, p = 0.729). These were exploratory analyses based on a very small sample however (n = 7 CH cages, with Cohort A represented by a single cage) and should be interpreted with caution.

## 5 Discussion

This chapter tested three hypotheses concerning housing quality, reproduction, and maternal care, developed in detail in the Introduction, Methods, and Results chapters: that housing quality (WR vs CH) shapes maternal and reproductive outcomes in DBA/2 dams; that it does so via one or both of two candidate mechanisms, namely lower BMI and/or elevated SB in CH females; and, if housing does affect reproduction, by which of these routes (or some other route) it does so. Rather than restate that reasoning here, the sections below take each hypothesis in turn and evaluate it directly against the results.

### 5.1 Housing affected some, but not all, reproductive and maternal outcomes

The first, basic hypothesis that housing quality (WR vs CH) shapes maternal and reproductive outcomes in DBA/2 dams was supported, with maternal care showing the clearest housing effects. WR dams nursed their pups significantly more than CH dams, and built significantly better nests with higher nest scores. Their pups also showed a trend towards being heavier at PND 28. All these measures point in the same direction: poorer housing is associated with reduced or lower-quality maternal investment, consistent with the hypothesis that housing shapes maternal care.

Housing quality did not, in contrast, affect the rate of successful mating or pregnancy, nor most litter-level outcomes: litter size, live-born pup number, survivability, and pup weight at birth, PND 16, or PND 21 all showed no significant housing difference between CH and WR mothers. This indicates that CH housing did not compromise basic capacities to conceive, carry, and provision a litter through most of the pre-weaning period. This is consistent with Carvalho et al. (2009), who found that housing condition did not significantly alter mating success, litter size, live births, pup survival, or litter weights across lactation (days 4–18) over five consecutive gestations in Swiss mice. It is also consistent with my own earlier work in DBA/2 mice, in which all trio-housed females mated regardless of housing (Anandarajan et al., 2026).

DBA/2 mice already have a modest reproductive baseline: even under typical conventional housing, the strain produces small litters (Charles River Laboratories, 1999; The Jackson Laboratory, 2024). In a classic cross-strain comparison, DBA/2J mice had smaller litters than C57BL/6J (5.4 vs. 7.0 pups per litter) and a lower overall fecundity index (15.8 vs. 23.5), and they responded poorly to superovulation, although their natural mating rate was not the lowest among common inbred strains (Silver, 1995). If CH housing had simply added another chronic stressor on top of this baseline (cf. Cait et al., 2022), we might have expected mating, pregnancy or litter-size deficits to appear specifically in CH mothers. We found none. CH housing therefore did not seem to push this strain’s fecundity below a further threshold, at least at our sample size.

This pattern also fits the neuroendocrine framework introduced earlier. Stress can suppress the hypothalamic–pituitary–gonadal (HPG) axis, impairing gonadotropin release and fertility (Rivier & Rivest, 1991; Whirledge & Cidlowski, 2010; Toufexis et al., 2014). However, these effects depend on how intense and long-lasting the stressor is. Pregnancy is also partly buffered, because the maternal hypothalamic–pituitary–adrenal (HPA) axis becomes less responsive to stress at this stage (Brunton et al., 2008). Lactation, by contrast, is the most energetically demanding phase of reproduction in small mammals (Speakman, 2008), and may leave mothers with less capacity to absorb additional stress. Conception and gestation may therefore be more robust to the moderate chronic stress of CH housing than lactation-stage maternal investment proved to be in this study.

Together, these results point to a housing effect that emerges late, accumulating or becoming detectable only as pups approach weaning, rather than a deficit present from birth. This timing fits a maternal-investment effect that compounds in late lactation. During this period, good nests help dams conserve energy that can be redirected into pup provisioning (Gaskill, Pritchett-Corning, et al., 2013). Consistent with this, nest score in the present study showed a trend towards predicting nursing, with higher nest scores associated with more nursing. This trend did not reach significance, and with this sample size it should be treated as suggestive rather than conclusive. WR dams also nursed for significantly more days before natural weaning than CH dams, indicating that CH dams weaned their pups earlier (although ‘walk-off’ behaviour, a separate index of natural weaning, did not differ by housing). Earlier weaning by CH dams could on its own explain a late-emerging weight gap, because pups depend heavily on milk until they begin eating solid food in the third postnatal week (König & Markl, 1987). It could also act alongside differences in nursing quantity or nest quality.

Pup retrieval, however, produced the most striking and unexpected result: CH females retrieved pups successfully in 100% of cages, while WR females did so in 0%, a complete separation between groups. Read alongside the poorer nursing, nest-score and PND 28 weight results of CH dams, this outcome appears to run counter both to the general pattern of poorer maternal care under CH housing and to evidence that chronic stress impairs pup retrieval (e.g., Abdelmesih et al., 2023). It should therefore be interpreted with caution rather than taken at face value as evidence that CH mothers retrieve pups more readily. I propose that the difference may reflect how dams from each housing type perceived "nest" versus "non-nest" space. For this study, all tests were carried out in each dam’s own home cage: the conventional cage for CH dams, and, for WR dams, the detachable (conventional) annex, which was identical in size and layout. WR dams built their nests mostly in this annex. Relative to the much larger WR cage as a whole, the annex may therefore have functioned as a dedicated nesting chamber, much as wild house mice nest in discrete chambers within a larger burrow and territory (Latham & Mason, 2004). If WR dams perceived the entire annex as nesting area, a pup displaced within it may not have been perceived as separated from the nest at all, and so would not have triggered retrieval. For CH dams, by contrast, the same-sized cage was their whole living space, within which the nest occupied only a small, clearly bounded area, so a displaced pup would have been outside the nest and in need of retrieval.

Other, non-exclusive explanations are also possible. CH dams, housed in conditions associated with greater anxiety and poorer welfare (Nip, 2018; Cait et al., 2022; MacLellan, 2023), may have perceived pup displacement as more threatening. Consistent with this, rat dams in CH cages have been reported to show more anxiety-like behaviour and higher corticosterone while spending *more* time with their pups (Cannervik, 2011). CH dams’ high levels of SB might also reflect broader behavioural disinhibition, which has been linked to cage stereotypies in bank voles (Garner & Mason, 2002), so that they responded to a displaced pup more impulsively. Alternatively, CH pups may have vocalised more readily when displaced. Pup ultrasonic calls reflect pups’ emotional state, increase with cooling, and are a key trigger of maternal retrieval (Branchi et al., 2001; Ehret, 2005; D’Amato et al., 2005). Given their dams’ poorer nests, CH pups may have cooled faster outside the nest (cf. Gaskill, Pritchett-Corning, et al., 2013). Whether CH and WR pups differed in calling or thermoregulation was not measured here. The retrieval result may therefore reflect differences in how the test was experienced across housing types, rather than reduced maternal responsiveness in WR dams.

### 5.2 Housing strongly affected SB but not BMI

The housing manipulation produced the expected, large effect on stereotypic behaviour. CH females showed significantly more stereotypic behaviour than WR females both before mating and on PND 17; the effect persisted from PND 17 to PND 24. This is consistent with the well-established sensitivity of barren, spatially restrictive housing to inducing stereotypic behaviour across taxa (Mason, 1991; Mason & Latham, 2004). It is also consistent with direct evidence that conventional housing induces stereotypic behaviours in mice, including DBA/2 mice (Nip, 2018), and with broader evidence that housing-induced neurobiological changes are implicated in stereotypic behaviour in laboratory mice (Kitchenham et al., 2024).

By contrast, housing had no significant effect on BMI at either time point (pre-mating and PND 16). This is a pivotal result for the energy hypothesis: for BMI to mediate a housing effect on reproductive or maternal outcomes, housing must first affect BMI, and it did not do so at either stage. This contrasts with MacLellan (2023), where conventionally housed DBA/2 females had lower BMIs than well-resourced conspecifics. However, those mice were assessed in late adulthood, after more than a year in their housing treatments, considerably longer than the dams in the present study. One possibility, therefore, is that our housing manipulation, while sufficient to produce a very large SB difference, was not prolonged or severe enough to produce a comparable shift in energy balance. This would also distinguish our design from studies reporting weight loss under more extreme SB-inducing conditions in horses (Christie et al., 2006) and mink (Jeppesen et al., 2004). Other, non-exclusive explanations are also plausible. Our dams had ad libitum access to food, which may have buffered any increase in energy expenditure from SB. The SB scored here may also have been dominated by low-energy, small-amplitude movements rather than the more energetically costly, whole-body locomotor stereotypies, such as route-tracing, most clearly linked to weight loss in the mink and horse literature.

### 5.3 Does SB predict reproduction and maternal care?

Having established that housing affected SB, we next asked whether SB explained the housing effects on reproduction and maternal care. If SB mediated these effects, two things should follow: adding SB to the model should weaken the housing effect, and SB itself should predict the outcome. Across outcomes, the first pattern appeared repeatedly, but the second did not reach significance for any outcome.

#### Nest score

Housing alone significantly predicted nest score, with WR dams building better nests. Once SB was added, the housing effect weakened to non-significance, and SB showed a trend towards predicting nest score, with higher-SB dams building poorer nests. This pattern is suggestive of partial mediation by SB, but it does not demonstrate it. The estimates are also unstable, because two of the four nest-score categories contained only a single observation each. With that caveat, the direction of the trend matches findings in farmed American mink, where dams showing more SB built poorer, flatter nests (Díez-León, 2014), and where enrichment that reduced SB also improved nest construction (Clark et al., 2025).

However, nest scores may also reflect the materials available. Mice are strongly motivated to obtain nesting material (Van de Weerd et al., 1998), and WR dams could carry additional materials, such as shredded paper, nesting squares and cardboard, from the wider enclosure into the detachable cage where they usually nested, whereas CH dams had only the two nesting materials common to both conditions. Because the detachable cage was identical to a CH cage, the nest-site enclosure itself is unlikely to explain the difference. However, with the present design, a direct effect of nesting resources cannot be separated from an SB-related or other maternal-state effect.

#### Pup weight at PND 28

Neither SB nor housing predicted pup weight at PND 28 once both were in the model. Because the housing effect on pup weight was only ever a trend, this is best read as a modest housing effect being diluted further, rather than a clear effect being explained away by SB. There was no sign that SB itself related to pup weight. The trend-level housing effect on pup weight therefore does not appear to be a downstream consequence of maternal SB, at least not detectably with this design and sample size. This contrasts with American mink, where high maternal SB predicted slower offspring growth (Díez-León & Mason, 2016). The difference may reflect species, the measures used, and/or limited statistical power here.

#### Nursing behaviour and number of nursing days

Nursing behaviour and total nursing days showed the same attenuation pattern as nest score, but without any SB trend. Housing alone significantly predicted both outcomes, but housing quality no longer did so once SB was included, and SB did not significantly predict either outcome. Some of the raw housing association with each measure can therefore be statistically accounted for by variation in SB. However, a genuine mediator would normally retain an association with the outcome after the exposure is accounted for, which SB did not. We therefore interpret this cautiously rather than as evidence of mediation. The pattern parallels findings in farmed mink, where an enrichment intervention that reduced SB and improved nest quality did not measurably improve nursing time (Clark et al., 2025). Weak or absent associations between SB and maternal care have also been reported in C58/J mice, a strain showing pronounced SB (Ryan et al., 2010).

#### Pup retrieval

Among CH cages, SB did not predict latency to retrieve the first pup or latency to bring all pups to the warmer area. With only seven cages, one cohort represented by a single cage, this is preliminary at best. Nevertheless, it indicates that CH dams did not retrieve pups faster because they were more stereotypic. Processes captured by SB, whether stress or impulsivity, are therefore unlikely to explain the between-groups retrieval pattern.

#### To conclude

Overall, adding SB to the models weakened housing effects on nest score, nursing time and nursing days, and SB showed a trend towards predicting nest score. However, SB did not significantly predict any outcome, including pup weight and pup retrieval latency. These results thus offer, at best, mixed and only partial support for SB as the mechanism linking housing to maternal care: the specific SB measure used here only partly accounts for housing’s effects. Our composite SB score may also have masked a real association. It combined behaviourally heterogeneous forms of SB into a single measure: higher-energy locomotor patterns (e.g., route-tracing) and lower-energy stationary patterns (e.g., bar-mouthing). If these subtypes relate differently to maternal care, combining them could dilute a true effect.

Some combination of a genuine but underpowered SB contribution, an unmeasured consequence of poor housing, or a more direct effect of housing quality most likely underlies the outcomes affected. This is consistent with evidence that housing and enrichment effects on affective outcomes do not always track individual differences in stereotypic behaviour (Gross et al., 2011; Kitchenham et al., 2024). It is also consistent with similar dissociations for maternal outcomes, mostly in mink, where enrichment reduced SB and improved nest building without clearly improving other maternal care outcomes (Clark et al., 2025; Díez-León & Mason, 2016).

### 5.4 Limitations

There are three main limitations that should be considered when interpreting these findings. First, this study directly tested only two of several plausible mechanistic pathways: the energy/BMI and ARB/SB hypotheses. Even for these two, food intake of dams and pups was not measured, which would have helped explain the absence of a BMI effect, and SB was recorded as a single category rather than by subtype.

Second, the sample size was small. Although there were N = 36 cages overall, this fell to as few as 7–8 cages per housing group in the maternal care analyses. This limits statistical power to detect small or moderate effects, as reflected in the small denominator degrees of freedom in several models. Results interpreted here as null should therefore be read as “not detected at this sample size” rather than as confirmed absences of effect. This applies particularly to the SB analyses, where the attenuation of housing effects without significant SB effects may well reflect limited power. With hindsight, DBA/2 mice may therefore not have been the best strain for these questions. They were chosen because they show high levels of SB and small litters under conventional housing, but their poor breeding limited sample sizes; a more fertile, outbred strain that also shows SB, such as CD-1 mice (Gross et al., 2012), would have allowed the SB pathway to be tested with greater power.

Finally, the pup retrieval assay may not have measured the same thing in both housing types. Retrieval was tested in each dam’s home cage, which for WR dams was the annex where they mostly nested. This cage was physically identical to the CH cage but may have functioned as a dedicated nesting chamber within a larger home range, so a displaced pup there may not have registered as separated from the nest. Pups displaced within this nesting area may also have called less, so pup ultrasonic calls should be recorded in any future replicate.

## 6 Conclusions

Taken together, these results support the hypothesis that housing quality shapes maternal and reproductive outcomes in DBA/2 dams, but they show that its effects are selective. Mating success, pregnancy and most litter-level measures were unaffected by housing. In contrast, CH dams nursed less, weaned their pups earlier and built poorer nests than WR dams, and their pups showed a trend towards lower weight at PND 28. Pup retrieval also differed between housing types, but in the opposite direction to that expected, which may have reflected differences in how the test was experienced across housing types rather than greater maternal responsiveness in CH dams. Housing therefore appears to have mainly affected how dams cared for their pups during lactation, and possibly pup growth as a result, rather than their ability to conceive and carry a litter.

How housing produced these effects remains unresolved. The energy hypothesis was not supported, as housing did not affect BMI. SB, although strongly increased by CH housing, showed only a trend towards predicting nest score and did not predict any other outcome, and housing itself remained the more consistent predictor. The weakening of housing effects once SB was included nonetheless leaves open a partial mediating role for SB that deserves better-powered testing. Other routes, including neuroendocrine changes, depression-like states, milk quality and nesting opportunity itself, could not be tested here and remain plausible; how all of these pathways could be tested is set out in Chapter 5.

These results have practical implications beyond DBA/2 mice. In laboratory mouse breeding facilities, maternal care directly affects colony productivity and animal welfare. Well-resourced housing of the kind tested here could plausibly improve nest quality and nursing, and possibly offspring weight around weaning, even without a confirmed mechanism and even where fertility and litter size are already adequate under standard housing. A more speculative possibility is that combining enrichment with a modestly delayed weaning age might compound this benefit. If enriched housing tends to produce heavier, better-developed pups by PND 28, delaying weaning slightly further, within welfare-appropriate limits, might allow pups to leave their dam in better condition, with possible knock-on effects on later-life traits such as reduced SB (Würbel & Stauffacher, 1998), enhanced male attractiveness to females (Anandarajan et al., 2026) and female reproductive performance. We are not aware of this combination having been tested directly, and offer it as a testable extension of the present findings rather than an established result.

## Ethics statement

We confirm the mate choice experiment was conducted with approval from the University of Guelph, Animal Care Committee (AUP #4570) and complied with Canadian Council of Animal Care guidelines.

## Acknowledgments

The authors acknowledge that this work was completed on the ancestral lands of the Attawandaron people and the treaty lands and territory of the Mississaugas of the Credit. The authors would also like to thank all the mice used in this study; all the adopters who gave our retired mice a caring home; our funder Natural Sciences and Engineering Research Council (NSERC), Canada, ACS Veterinarian Dr.Sarah Baert, Animal Care Technicians Michaela Randall, Tammy Buitenhuis and Michelle Cieplak, past Mason Lab Master’s students Lauren Young, Shay Forget and Jiayi Cai, as well as Research Assistants Ainsley Robertson and Katrina Seniuk.

## Disclosure of AI use

We acknowledge that Prathipa Anandarajan used AI-assisted tools (Grammarly and Claude) to assist with grammar, readability, clarity, and rephrasing of her own sentences without changing their meaning. These tools were not used to generate ideas, arguments, interpretations, or other substantive intellectual content. All conceptual contributions and conclusions were developed solely by the authors. No confidential information was provided to these tools.

